# Nanoscaled discovery of a shunt rifamycin from *Salinispora arenicola* using a three-colour GFP-tagged *Staphylococcus aureus* macrophage infection assay

**DOI:** 10.1101/2022.12.04.519019

**Authors:** Nhan T. Pham, Joana Alves, Fiona A. Sargison, Reiko Cullum, Jan Wildenhain, William Fenical, Mark S. Butler, David A. Mead, Brendan M. Duggan, J. Ross Fitzgerald, James J. La Clair, Manfred Auer

## Abstract

Antimicrobial resistance has emerged as an urgent global public health threat, and development of novel therapeutics for treating infections caused by multi-drug resistant bacteria is urgent. *Staphylococcus aureus* is a major human and animal pathogen, responsible for high levels of morbidity and mortality worldwide. The intracellular survival of *S. aureus* in macrophages contributes to immune evasion, dissemination, and resilience to antibiotic treatment. Here, we present a confocal fluorescence imaging assay for monitoring macrophage infection by GFP-tagged *Staphylococcus aureus* as a front-line tool to identify antibiotic leads. The assay was employed in combination with nanoscaled chemical analyses to facilitate the discovery of a novel, active rifamycin analogue. Our findings indicate a promising new approach to the identification of anti-microbial compounds with macrophage intracellular activity. The novel antibiotic identified here may represent a useful addition to our armoury in tackling the silent pandemic of antimicrobial resistance.

---

Traditional targeted high-throughput screening (HTS) of large chemical libraries created by combinatorial chemistry has proved very efficient in finding hits and lead compounds for a variety of indications. It was thought that this process would be similarly successful in discovering antibiotics. However, several attempts to use HTS against large corporate combinatorial libraries have failed to identify new broad-spectrum antibiotics^1–3^. Furthermore, common antimicrobial screening methods that analyse bacterial growth *via* target-based biochemical assays can overlook possible cytotoxic effects and permeability to cellular membranes^4^. For these reasons, allied to the automation and development of high content screening (HCS) technologies, drug discovery strategies have turned to cell-based assays that employ the microbe in its cellular environment^5^. In 2009, Christophe *et al.* employed HCS to identify synthetic chemical compounds to inhibit the intracellular replication of *Mycobacterium tuberculosis* within macrophages^6^. Subsequent studies led to the identification of a potent clinical candidate, telacebec (Q203), for the treatment of tuberculosis (Phase II study completed) as well as for Buruli-ulcer (Phase I study underway in Korea)^7^. Related assays have been used to identify antimicrobial compounds against *Salmonella typhimirium*^8^, *Plasmodium falciparum* and *Leishmania* spp.^9^. These assays are especially important for the discovery of antimicrobial agents active against intracellular pathogens or pathogens that survive and replicate inside host cells during their infection cycle^5^.

*S. aureus* is a leading cause of hospital and community acquired infections and its ability to develop resistance to antimicrobial agents makes it a priority for the development of agents with new modes of action, superior toxicity profiles, and intravenous (IV)/oral switch administration. During *S. aureus* pathogenesis, the interaction with host macrophages has a pivotal role in determining the outcome of infection. *S. aureus* has evolved multiple strategies to survive within, manipulate, and escape from macrophages^10^. The macrophage intracellular environment not only shields the bacteria from most antimicrobials, but also can support the development of persister cells with an antimicrobial tolerant phenotype^11, 12^. To address these issues, we developed a high content human macrophage infection assay (Fig. 1a) to identify new antibiotics from marine microbial test fractions with antimicrobial properties against multi-drug-resistant *S. aureus*. With this assay we identified intracellular *Staphylococcus aureus* growth inhibitors in in macrophages^13^.

**Fig. 1.**
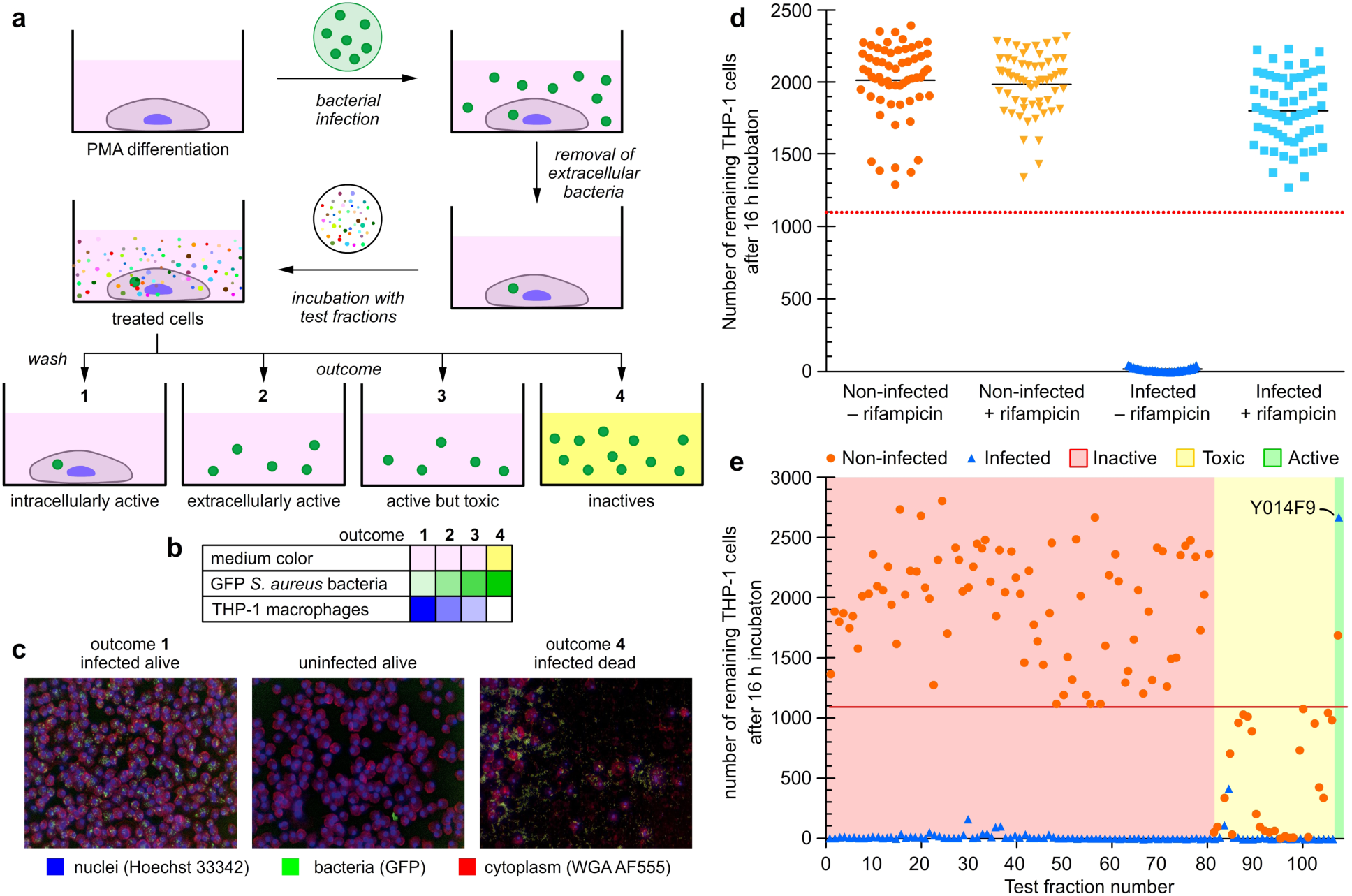
A fluorescent macrophage infection assay. **a** Schematic representation of the three-colour confocal fluorescence macrophage infection assay run in 96-well plate format on a Perkin Elmer Opera instrument using a 20× objective. The assay process begins with infecting PMA differentiated THP-1-macrophage cells with GFP-USA300 *S. aureus*. Co-culturing of these cells results in infected mammalian cells after washing and removal of the media. The resulting bacterial infected cells are then subjected to test samples, which can result in four different outcomes. The first, arising from intracellular active compounds, is identified by a significant number of infected THP-1 cells remaining after the overnight incubation (indicated as dark blue in Fig. 1b) and the cell media is red/pink (indicated as rose in panel b). The number of GFP- USA300 *S. aureus* bacteria is low (indicated as light green in panel b). The next two, extracellular active and active but toxic compounds, are marked by a low number of remaining THP-1 cells (indicated as light blue in panel b), some extracellular bacteria (indicated as lighter green in Fig. 1 b, and red/pink cell media (indicated as rose in panel b). The only difference between these two outcomes is that in the non-infected control only a few to no THP-1 cells are found for toxic compounds. The final, arising from non-active compounds, contains an absence of THP-1 cells as well as yellow cell media (panel b, lane 4), caused by saturated growth of bacteria (indicated by a dark green coloured field in panel b, lane 4), which turns the cell media acidic, causing a colour change to yellow. The cell membranes and the nuclei of the THP-1 macrophages were stained with wheat germ agglutinin Alexa Fluor 555 (WGA AF555) and Hoechst 33342 to enable blue/green/orange detection scheme. **b** A heatmap showing the outcome for each of the four different possibilities. The colour in the table illustrate whether there is a colour change in the cell medium and the strength of the colour illustrates the number of bacteria or THP-1 cells found in the sample. Medium colour: colour change from pink (red) to yellow only for outcome 4. GFP *S. aureus* bacteria (green) are most abundant for outcome 4. THP-1 macrophages (blue) are most abundant for outcome 1 and absent for outcome 4. **c** Typical Opera images for the various outcomes of the assay, THP-1 nuclei and plasma membrane are blue and red, respectively and *S. aureus* bacteria are in green. Left image visualizes the outcome for an intracellularly active compound, the middle for an extracellularly active compound and right image for an inactive compound. **d** Assay sensitivity. The separation between positive control containing infected THP-1 cells in 50 nM rifampicin (light blue square) and negative control (dark blue upright triangle) containing infected THP-1 cells in media only. For comparison non-infected THP-1 cells with (yellow upside down triangle) and without (orange round dots) 50 nM rifampicin are also shown. The red dashed line is the hit threshold which is the lower 3σ (standard deviation) or limit of the average of infected THP-1 cells in 50 nM rifampicin (parameters providing in the Supplementary Information). **e** Results of the screen of 108 marine microbial test fractions. More than 80 fractions were inactive (red shaded area) and the rest of the fractions were toxic to THP-1 cells (yellow shaded area) except fractionY014F9, which was identified to be active, with a high number of infected and non-infected THP-1 cells remaining after overnight incubation.

## Results

### Design and implementation of a three-colour GFP-tagged Staphylococcus aureus macrophage infection assay

To address the failure to identify new antibiotics from drug-like small molecule chemical libraries using conventional screening methods, we developed a new *S. aureus* macrophage infection assay. Using this assay, we screened 108 marine natural product fractions, consisting of ∼1080 compounds (average of 10 compounds per fraction), for the ability to inhibit intracellular *S. aureus* growth and survival inside macrophages. Our assay (Fig. 1a) utilised THP-1-derived macrophages infected with a green fluorescence protein (GFP) expressing a community-associated, methicillin-resistant *S. aureus* (MRSA) strain (USA300 LAC-GFP)^14^. THP-1 cells are a human monocytic cell line that are differentiated into macrophages using phorbol myristate acetate (PMA). These cells present an inflammatory macrophage phenotype, are highly phagocytic^15^ and have been extensively used to study bacterial-macrophage interactions^16^ and antimicrobial intracellular activity^15, 17^. Despite some differences when compared with primary cells^18^, the use of this cell line allowed us to overcome issues with availability, donor variability and ethical considerations associated with primary human blood monocyte-derived cultures, as well as high reagent costs and the time consuming protocols of preparing macrophage-like cells from human embryonic stem cells^19^. The differentiated, adherent THP-1 cells were infected with USA300 LAC-GFP *S. aureus* at a multiplicity of infection (MOI) of 10 for 1h and, in order to test the intracellular activity of the compounds, the extracellular bacteria were killed by incubation with gentamicin prior to the addition of the test fractions.

Natural product fractions were added to the cells at 25 µg/mL and 5 µg/mL and incubated for 16 h. Subsequently, cells were fixed with 4% paraformaldehyde (PFA) and stained with Hoechst 33342 (nuclei) and Wheat Germ Agglutinin Alexa Fluor 555 Conjugate (plasma membrane) giving a three-colour read out with *S. aureus* stably expressing GFP. Using this assay, we observed four possible outcomes: inactive, active but cytotoxic, extracellular active, and intracellular active extracts (Fig. 1a-b). As in the negative control (media only), incubation with inactive compounds allowed the bacteria to kill the THP-1 macrophages, escape the intracellular environment, and multiply in the cell culture media during overnight incubation, lowering the pH of the media facilitated its colour change from red to yellow. Extracellularly-active fractions had no colour change of the media, which was due to absence of both saturated bacterial growth and live THP-1 macrophages. We focused on identifying intracellular active compounds associated with no media colour change and the presence of live THP-1 cells after the overnight incubation (further details in the Methods). As shown in Fig. 1d, we were able to validate our assay using rifampicin as an active model drug. Over multiple repetitions of the imaging process, infected THP-1 cell death occurred within 16 h, which was prevented in cultures treated with rifampicin (Fig. 1d).

### Screening of marine microbial test fractions leads to a hit fraction Y014F9

Next, we turned to chromatographically-fractionated marine microbial fractions (test fractions) developed in the Fenical laboratory from a large repository of marine microbes (> 17,000 strains)^20^. A panel of 108 test fractions was selected that contained ∼10 compounds per fraction at a concentration of 10-50 mg/mL in DMSO. Here the goal was to complete the entire discovery effort with less than 1 mL of each test fraction and finish with a characterized active compound. To achieve this, we needed to determine the limits of this screening effort and began by screening 100 µL aliquots of these extracts. For each extract, a 500 µL aliquot was saved for compound purification. Fig. 1e shows the outcome of the primary screen of the 108 extract fractions. About a quarter of the fractions showed toxicity (at 25 µg/mL) against macrophages. This can be seen by the low number of THP-1 macrophages in the non-infected controls, indicated by the orange data points below the red line in the yellow shaded part of Fig. 1e. There were no remaining infected THP-1 macrophages for all but a few extracts. However, only extract Y014F9 yielded macrophage cell numbers exceeding our hit criteria (above the red line, detailed description of the assay parameters and hit threshold is given in the Methods), blue data point marked with Y014F9 in Fig. 1e.

### Compound identification and validation through microbial reculturing

With a hit test fraction (Y014F9) identified, we turned to methods scaled at the nanomolar level (nanoscaled) to conduct the assay-guided fractionation directly from the screening aliquot. A sample of the DMSO stock of test sample Y014F9 (100 µL) was dried by airflow and redissolved in MeOH (50 µL). A 50 µL solution of the MeOH stock was separated using a 5 cm × 20 cm, pTLC plate (250 µm) (eluent 1:9 mixture of MeOH:EtOAc) into eight bands (Y014F9-1 to Y014F9-8). Each band was submitted to nanoscaled-NMR analyses. Half of the remaining material was resubmitted to the assay, while the other half was saved for further purification. Application of the assay to the fractions identified Y014F9-6 (NMR spectrum provided in Fig. 2a) as the most active. Using the remaining half of the material, we repeated the fractionation to afford three bands Y014F9-6A to Y014F9-6C (NMR spectra provided in Fig. 2a). After collection of NMR data, these samples were then resubjected to the assay identifying Y014F6-6B as the most active (Fig. 2b).

**Fig. 2.**
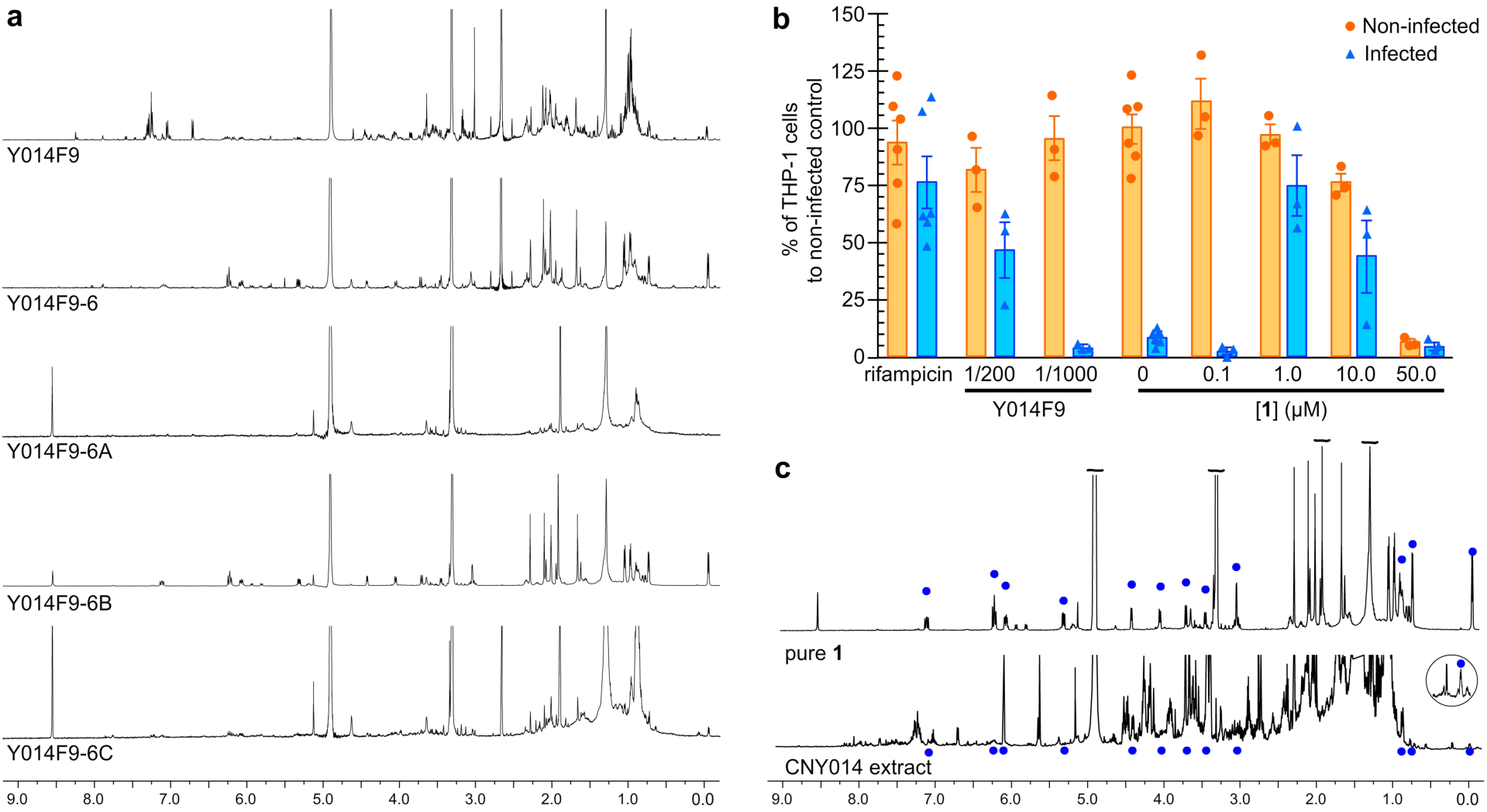
Nanomole-scaled isolation of 1 from the test fraction Y014F9. **a** ^1^H-NMR spectra collected on 50 µL sample of the test fraction Y014F9. A 100 µL sample of the test fraction in DMSO was then further fractionated into 8 bands by pTLC to afford fractions Y014F9-1 to Y014F9-8. A second purification of Y014F9-6 afforded three cuts Y014F9-6A to Y014F9-6C. **b** Testing of compound **1** obtained from Y014F9 at 0.1, 1.0, 10 or 50 µM in the THP-1 macrophage infection assay. For comparison, 50 nM rifampicin and the parent test fraction Y014F9 at 1/200 and 1/1000 dilution are shown. **c** ^1^-NMR spectra comparing purified **1** against the CYN014 extract.

We then recultured the microbe, *Salinospora arenicola* strain CNY-014, that had produced test fraction Y014F9. Repetition of the culturing at 5 L scale returned 400 mg of crude extract, which after NMR guided-fractionation afforded 8 mg of fraction S10 (Supplementary Table 1). Further purification of this material by prep-TLC (5 cm × 20 cm, 250 µm pTLC plate and elution with 1:9 mixture of MeOH:EtOAc) afforded 520 µg (estimated by NMR) of rifamycin analogue **1** (Fig. 3a). By ^1^H-NMR analysis (Fig. 2c), this material was identical to that obtained from purification of Y014F9-6B (Fig. 2a), thereby confirming the source as well as providing sufficient material for NMR analyses. That achieved, the low yield (0.13%) of **1** indicated that it was a minor component of the extract, a fact that was clearly evident by comparing the NMR spectrum of pure **1** to its parent extract (Fig. 2c).

**Fig. 3.**
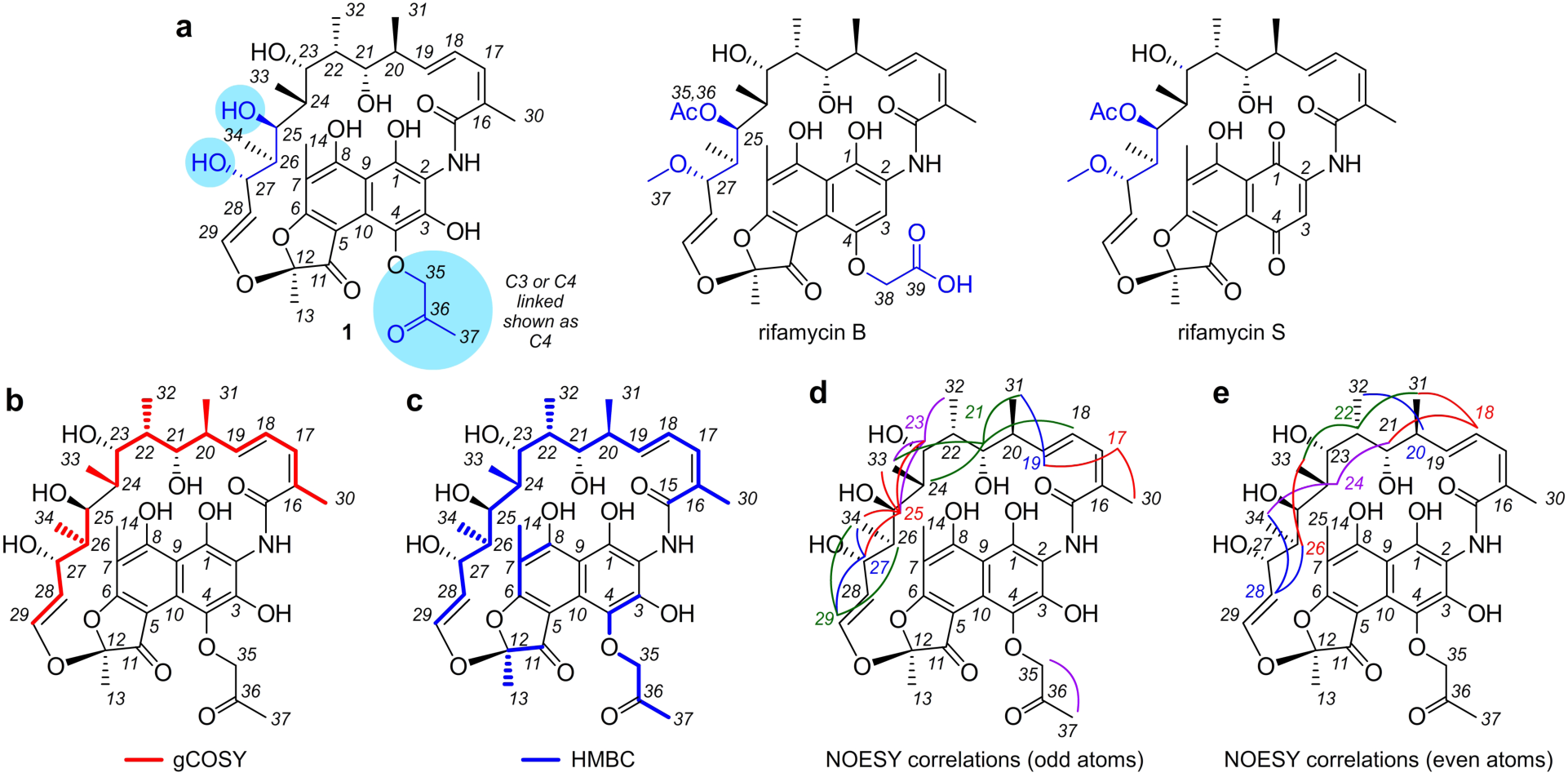
Structure elucidation. **a** Structures of 4-(propan-2-one)-25-*O*-deacetyl-27-*O*- desmethylrifamycin **(1)**, rifamycin B and rifamycin S. Reference NMR datasets were collected from rifamycin B (Supplementary Table S3) and rifamycin S (Supplementary Table S4) to aid in the structure elucidation process. **b** A summary of the ^1^H-^1^H gCOSY data obtained from **1** in CD_3_OD (Supplementary Table S2). Key gCOSY (red) correlations are shown on **1**. **c** A summary of the ^1^H-^13^C HMBC data obtained from **1** in CD_3_OD (Supplementary Table S2). Key HMBC (blue) correlations are shown mapped on **1**. **d,e** The stereochemical assignments in **1** were confirmed by tabulating the ^1^H-^1^H NOESY data from **1** (Supplementary Table S2) and comparing them to that expected from X-ray crystal structural data from rifampicin (Supplementary Fig. 3). Key NOE interactions are displayed in **d** for the odd numbered protons from H17-H29 and **e** even numbered protons from H18-H28. Search engines including Scifinder, MarinLit, and Coconut indicate that **1** is a novel member of the rifamycin class. While 25-*O*- deacetyl-27-*O*-desmethylrifamycin is a known motif, all evidence from these searches suggest that this is the first member of this large family of natural products with a propan-2-one modification.

### Structure elucidation and activity validation of the active component

With 520 µg of **1**, we were able to obtain a dataset including ^1^H-^1^H gCOSY, ^1^H-^13^C HSQC, ^1^H-^13^C HMBC and ^1^H-^1^H NOESY and HRMS spectra. HRMS analyses returned an *m/z* [M-H^-^] of 713.3042, which was consistent with the formula of C_37_H_47_NO_13_ (calcd. 713.3047). While database searching (MarinLit, SciFinder) failed to produce hits, we were rapidly able to build two fragments by interpretation of ^1^H-^1^H gCOSY NMR data (red lines, Fig. 3b) along with ^1^H and ^13^C assignments from the ^1^H-^13^C-HSQC spectrum. ^1^H-^13^C HMBC NMR data were then used to identify the missing correlation between C24-C25 in the gCOSY data resulting in a single 20 carbon fragment (blue line, Fig. 3c).

Based on prior studies in *Salinospora*,^21^ we found that resonances within the NMR spectra from **1** matched those observed in the ansamycin family of polyketides^22^. In particular, the presence of three olefinic protons coupled within a diene (confirmed by ^1^H,^1^H-gCOSY and ^1^H,^13^C-HMBC data), four oxygen substituted methines and 4 methyl groups, also methine-attached, strongly supported this conclusion. Furthermore, the ^1^H-^1^H gCOSY and ^1^H-^13^C HMBC data showed additional correlations to a disubstituted olefin with a ^3^*J*_HH_ coupling constant of 12.7 Hz, all of which indicated that **1** was a new member of the rifamycin subclass^23^ of the ansamycin family.

We then collected NMR data (^1^H-^1^H gCOSY, ^1^H-^13^C HSQC, ^1^H-^13^C HMBC and ^1^H-^1^H NOESY) from rifamycin B (Fig. 3a) and rifamycin S (Fig. 3a) as tools to aid structure elucidation. First, we determined that the C28-C29 olefin in **1** contained comparable proton coupling constants (^3^*J*_27-28_ of 7.0 Hz, ^3^*J*_28-29_ of 12.7 Hz and ^4^*J*_27-29_ of 1.4 Hz, Supplementary Table 2) to those observed in rifamycin B (^3^*J*_27-28_ 7.0 Hz, ^3^*J*_28-29_ 12.7 Hz and ^4^*J*_27-29_ 1.0 Hz, Supplementary Table 3) and rifamycin S (^3^*J*_27-28_ 7.9 Hz, ^3^*J*_28-29_ 12.6 Hz and ^4^*J*_27-29_ 0.8 Hz, Supplementary Table 4), therein indicating that the olefin residue observed by ^1^H-^1^H gCOSY (Fig. 3b) and ^1^H-^13^C HMBC (Fig. 3c) analyses was likely a *trans*-olefin, as observed in rifamycins B and S.

Chemical shift analyses (^1^H and ^13^C, Supplementary Fig. 1) were used to identify the functionality in **1** that was different from that in rifamycins B or S. Overall the chemical shifts of **1** shared a strong similarity with selected shifts from rifamycin B and rifamycin S (Supplementary Table 5); however, the aromatic proton H3 was not present (7.37 bs in rifamycin B and 7.63 s in rifamycin S, Supplementary Table 5), which indicated functionalization (commonly observed as oxidation) at C3. Functional modification between C25-C28 was suggested by chemical shift differences observed for H25, H27, C25, C27 and C28. Using this data as a guide, we were able to determine that **1** had secondary alcohols at positions C25 and C27, was not acetylated at C-25 as observed in rifamycin B (C35 δ172.6, H36 δ 2.02, C26 δ 20.8, Supplementary Table 3), and was not methylated at position C27 (H37 δ 3.03, C37 δ 57.1, Supplementary Table 3). In addition to these structural differences (blue highlights, Fig. 3a), the spectral data from **1** (Supplementary Table 5) did not contain the same glycolic acid on C4 found in rifamycin B, as noted by the lack of peaks associated with rifamycin B (H38 δ 4.74 and C38 δ 67.7 ppm). In contrast, **1** contained a 2-propanone functionality whose position was tentatively assigned to C4 (see note in Fig. 3a). Unfortunately, complete assignment of this position was complicated by the fact that in **1** the C15-C29 section of the molecule had multiple conformations as clearly evident by the presence of multiple peaks for the methyl singlets of C14, C30 and doublets of C31-C34. This was further supported by the presence of cross peak duplication in the ^1^H-^13^C HMBC spectrum and exchange peaks in the ^1^H-^1^H NOESY spectrum suggesting that at least two conformations were present.

While the polyketide synthase (PKS) derived region (C15-C19 and C30-C34) remained uniform, we soon discovered that the alkoxy-2-propanone (C35-C37 in **1**, Fig. 3a) was unstable and led to the formation of traces of **2** (Supplementary Fig. 2). This observation was further supported by the increase in the hydrolysis of **1** to **2** upon storage in CD_3_OD solution during NMR studies (data not shown). Ultimately, this added level of complexity led to formation of multiple sets of ^13^C peaks for the aromatic residues C1-C10 (ND in Supplementary Table 2). While methods were established to enable their assignment for rifamycin B (Supplementary Table 3) and rifamycin S (Supplementary Table 4), complete assignment of the aromatic resonances of **1** was not possible. The data reported in Supplementary Tables 3-4 were collected from samples of rifamycin B or S at comparable concentrations as **1** in CD_3_OD (note the inclusion of ^13^C NMR spectra and peaks within the aryl region of the ^1^H-^13^C HMBC for these two compounds).

With the carbon scaffold identified, we turned our attention to evaluate the stereochemistry of **1** from interpretation of ^1^H-^1^H NOESY NMR data (Fig. 3d-3e, Supplementary Table 2). Fortunately, a wealth of X-ray crystal structures of rifamycin analogues exists in the CDCC database. Using the crystal structure of rifampicin methanol solvate trihydrate (deposition CDC 859793), we systematically compared the observed NOEs (Supplementary Fig. 3) and coupling constants (Supplementary Fig. 4) with the configuration observed in the crystal structure data (Fig. 3d-3e) and found them consistent with a few exceptions. First, in solution, rotation was freely observed around the C27-C28 and C29-O bonds allowing the olefinic protons at C28 and C29 to interact with protons on both sides of the molecule. Similar rotation was observed between the diene from C16-C19. Nevertheless, the NOESY data not only confirmed the stereochemical assignment of the PKS portion of the molecule but also confirmed the full stereostructure assigned to **1**.

## Discussion

*S. aureus* is a leading cause of global mortality due to the increase in antibiotic resistance strains^24^, and new antibiotics are urgently needed. Using our new 3-colour THP-1 macrophage fluorescence imaging assay for intracellular *S. aureus* inhibition screening in iterative combination with nanomole-scaled techniques, we were able to identify a rifamycin analogue **1** that has not yet been described. The structure of **1** was shown to have a unique and readily activated alkyoxypropan-2-one motif. Samples of purified **1** were tested at a range of concentrations in our infection assay against *S. aureus* infected and non-infected differentiated THP-1 cells and the results are shown in Fig. 2b. The cell count of non-infected THP-1 cells suggested that although compound **1** was non-toxic at lower concentrations, toxicity was detected at 50 µM. Furthermore, cell count for infected cells indicates that **1** is most active at 1 and 10 µM. Combining activity and toxicity suggest that compound **1** has a narrow activity window of > 1 µM and < 50 µM. Using the Microplate Alamar Blue Assay (MABA) and Low Oxygen Recovery Assay (LORA)^25^ **1** demonstrated activity against *Mycobacterium tuberculosis* with MIC values of 1.3 µM and 16.8 µM, respectively. Parallel MABA analyses indicated that this compares favourably to rifamycin O (0.093 µM LORA), rifamycin SV (0.044 µM LORA) and rifampicin (0.13 µM LORA or 0.024 µM MABA). While active, the reduced activity of **1** is not unexpected given its need for oxidative activation to **2** (Supplementary Fig. 2).

## Conclusions

With the first three-colour intra-macrophage infection assay for *S. aureus* developed and demonstrated the stage is set for larger high-throughput screens for small molecular libraries and for the identification of novel natural products from extract screening. With the ultimate need to develop new antibacterial drugs to tackle the global AMR crisis, we expect that this protocol will be adapted for general use as it allows the identification of compounds that can target intracellular bacteria.

Marine actinobacteria are known for their prolific production of secondary metabolites^26^, many of which have demonstrated clinical utility^27^. Access to sequence data and new genomic tools has revolutionized our ability to analyse and manipulate large complex biosynthetic gene clusters (BGCs)^28^. This capability has advanced to the point where it is now possible to refactor old drugs for better treatments of infectious diseases^29^. Here, we apply a new screening approach to an unexplored, marine rifamycin producer, *Salinispora arenicola*^30^, to gain access to a new class of rifamycin analogues^31, 32^. On-going sequencing efforts illustrate the complex divergence within the biosynthesis of rifamycin (Supplementary Fig. 5c) from that observed in strains of *Amycolatopsis* and *Salinispora*. While the PKS portion of this pathway retains a common overall gene architecture (Supplementary Fig. 5a), a high degree of divergence is apparent within the pre-PKS supply (Supplementary Fig. 5b) and post-PKS tailoring enzymes (Supplementary Fig. 5d). Successful capture of these modifications through comparative genomic analyses provides an excellent opportunity to blueprint a minimal rifamycin synthase for synthetic biological applications. Here, the tremendous wealth of *Salinispora* genomes provides an excellent foundation to launch such efforts^33^.

The rifamycin group of antibiotics has demonstrated efficacy against bacterial and mycobacterial infections marked by its critical applications in the treatment of tuberculosis^34^, leprosy^35^ and Traveller’s diarrhoea^36^. Rifamycin polyketides are one of the few drug classes active against replicating and non-replicating bacteria, and new leads are critical to further clinical improvements. A recent survey of a series of strains of *S. arenicola* (Supplementary Fig. 6) indicated that extracts from ten selected strains produced rifamycins, whose structure and functionality has yet to be explored. Here, we demonstrate how the combination of a new assay and miniaturized isolation methods can effectively direct identification of new natural product as starting points for development of new drugs.

## Methods

### Preparation of THP-1 macrophages

THP-1 cells (kind gift from Prof. David Hume) were maintained in RPMI-1640 medium (Thermo Fisher) supplemented with 10% heat inactivated fetal bovine serum (FBS, Life Technologies) and GlutaMax (Thermo Fisher) at 37 °C in a 5% CO_2_ environment. Preparation of THP-1 cells prior to infection, THP-1 monocytes were differentiated into macrophages by treatment with 50 nM phorbol 12-myristate 13-acetate (PMA, VWR) for 3 days at a density of 5 × 10^4^ cells per well in a 96-well tissue culture plate (Thermo Fisher, Nunclon delta). The media was then replaced with fresh PMA-free media and the cells were left to rest for 24 h before the infection protocol. Cell differentiation following PMA stimulation was monitored over time by microscopic observation of cell morphology and plastic adherence properties.

### Macrophage infection with GFP-*S. aureus*

The *S. aureus* USA300 LAC strain that constitutively expresses green fluorescence protein (GFP) under a sarA-P1 promoter inserted markerless in the bacterial chromosome^14^ was used in this study. Bacterial overnight culture in tryptone soy broth (TSB, Oxoid) was sub-cultured in fresh TSB (1:100 dilution) at 37 °C, 200 rpm until reaching an OD_600_ value of 0.6. The bacteria solution was pelleted, washed and suspended in macrophage culture media to a concentration of 5 × 10^6^ bacteria/mL. A 100 µL aliquot of the bacterial solution was added to the THP-1 macrophage culture and the plate was centrifuged at 300 × g for 5 min to facilitate the macrophage-bacteria interaction. The infection was maintained for 1 h at 37 °C in a 5% CO_2_ atmosphere. Afterwards the media was replaced with media containing 100 µg/mL of gentamicin (Sigma, G1397) and left to incubate for 30 min to kill extracellular bacteria. The supernatant was then removed and the cells were washed twice with fresh media prior to addition of the test fractions.

### Addition of test fractions

A total of 108 marine microbial test fractions containing ∼5 mg/mL of dry extract in DMSO (Sigma Aldrich) were screened. For the screen the extracts were diluted in macrophage culture media at 200× and 1000× (corresponding to 0.025 mg/mL and 0.005 mg/mL, respectively) and added to the thoroughly washed THP-1 macrophages at total volume of 200 µl/well and the plate was left in the incubator overnight (16 h). Equivalent amount of DMSO was added to the cells in the DMSO control wells. Comparable methods were applied to test fractions and purified samples of **1**.

### Cell fixation and staining

THP-1 macrophages were then fixed with 4 % formaldehyde (formaldehyde 16% MeOH free, ThermoFisher Scientific) in phosphate buffered saline (PBS) for 20 min at 23 °C and subsequently washed thoroughly in PBS. Next the cell membranes and the nuclei of the THP-1 macrophages were stained with Alexa Fluor 555 conjugated wheat germ agglutinin (ThermoFisher Scientific) and Hoechst 33342 (ThermoFisher Scientific) in PBS, according to the supplier’s protocols. The cells were then washed twice thoroughly in PBS and a final volume of 100 µL of PBS was added to each well and the plate was stored at 4 °C until imaging.

### Imaging assays

Imaging was performed using the Opera HCS (Perkin Elmer) instrument. Two independent exposures with two fluorescence channels each were used to avoid bleed through. Exposure 1 parameters: laser excitation 488 nm at 2.5 mW, emission filter 520/35 nm. Exposure 2 parameters: laser excitation 561 nm at 2 mW and emission filter 585/40 nm and UV excitation at 365 nm and emission filter 450/50 nm, respectively. The exposure time for the three laser channels was 240 ms and the UV exposure time was 20 ms. All imaging was performed using a 20× Air LUCPLFLN objective (Olympus) with a NA of 0.45 and 20 images per well were captured.

### Image data analyses

Image analysis was completed using the image analysis software Acapella (Perkin Elmer). Two versions of the script were developed, one that is compatible with the Opera instrument to run while imaging (version 2.7) and one supporting instruments with Harmony and Windows 10 (version 5.2+). The script was calibrated using different treatment conditions: positive (50 nM rifampicin) and negative control (no rifampicin) treated wells, of *S. aureus* infected THP-1 macrophages and non-infected cells (Fig. 1c). The workflow is described in the Algorithm flowchart and example images are shown as well (Supplementary Fig. 7). Briefly, nuclei were detected using the algorithm H in the Hoechst channel, border objects were removed to ensure only whole nuclei were analysed. Area, roundness and intensity were calculated to filter objects downstream of the analysis. The cytoplasm was detected using algorithm B in the macrophage marker (cytoplasm channel). Those areas were kept as regions of interest to define individual cell areas. For those regions properties such as area and intensity and their standard deviations were calculated. A “find spots” algorithm was utilized for the infectious bacteria (*S. aureu*s GFP channel) foci analysis. Staining intensity and morphological properties were calculated in all channels and infectious ratio (normal cell fragmentation) was calculated for the cells identified (Supplementary Table 6). Potential artifacts in the analysis were removed using cell-like property filters. The remaining cell-like objects were transferred into a CSV output file. Final data analysis and graph plotting were performed in Excel and Prism (GraphPad).

### Development of assay parameters

Initially, the assay was tested with a range of mode of multiplicity to find the optimum. At MOI of 5, and below, the outcome of the assay was random. For example, when infecting cells in 60 wells at MOI 5, a significant number of wells show a large number of live macrophages remaining after the overnight incubation (without any addition of a test fraction or purified **1**) whereas in the remaining wells there were no live macrophages. Hence, the number of remaining live macrophages followed a stochastic distribution making the assay not useful. Therefore, we chose MOI of 10 to obtain a stable assay. To further characterise the assay *S. aureus* infected and non-infected differentiated THP-1 cells were kept overnight in complete RPMI-1640 medium with and without 50 nM rifampicin (Fig. 1d). After washing, fixating, staining, imaging and image analysis to identify the number of remaining THP-1 cells, dead cells were washed away during the fixing and staining procedures. These results are shown in Fig. 1. For non-infected THP-1 cells in the presence or absence of 50 nM rifampicin, there are a significant number of cells left after the incubation (Fig. 1d). For infected THP-1 cells in the absence of rifampicin (50 nM), cell death was predominant as noted by the presence of only a few cells after washing and fixation. However, in the presence of 50 nM rifampicin, there were a significant number of infected cells left, comparable to the numbers for the non-infected cells. This shows that our assay was sensitive with a large separation between our positive control (50 nM rifampicin) and negative control (cell media). However, it was difficult to use the negative control, due to the number of remaining cells being very close to zero, to define the threshold for hit identification. Hence, we chose the lower three-sigma threshold (three standard deviation) or a Z-score > −3 of the average cell number of the infected THP-1 cells in the presence of 50 nM rifampicin (red line in Fig. 1d) as our hit threshold, the minimum cell number a test fraction needed to be classified as a hit.

### Assay guided purification of 1

A 100 µL sample of the test fraction in DMSO was dried by air flow, redissolved in MeOH (50 µL), applied to a pTLC plate (5 cm × 20 cm), and eluted with a 1:9 mixture of MeOH:EtOAc. Eight bands (Y014F9-1 to Y014F9-8) were obtained and submitted to NMR analyses. Each of these fractions was rescreened to identify Y014F9-6 as the active fraction. ^1^H-NMR spectra were collected on a sample of each fraction. The remaining Y014F9-6 fraction was then redissolved in MeOH (50 µl) and applied to a pTLC plate (5 cm × 20 cm) and eluted with a 1:9 mixture of MeOH:EtOAc to deliver three fractions Y014F9-6A, Y014F9-6B, Y014F9-6C. A second repetition assay identified Y014F9-6B as the most active fraction.^1^H-NMR spectra were collected on a sample of each fraction. Comparable methods were used to screen the activity of purified **1**.

### Reculturing and fractionation of *S. arenicola* CNY-014

*S. arenicola* CNY-014 strain was cultured in 5 L scale for 7 days with shaking at 180 RPM. After this period the cultures were extracted with EtOAc (2 × 5 L), the extract dried with Na_2_SO_4_ and concentrated by rotary evaporation to yield 400 mg of organic extract. The ethyl acetate extract (400 mg) was subjected to silica chromatography, using a step gradient solvent system of n-hexane, EtOAc, and MeOH (1:0:0, 4:1:0, 3:2:0, 1:1:0, 1:4:0, 0:1:0, 0:9:1, 0:1:1, and 0:0:1) to yield 10 fractions as provided in Supplementary Table 1.

### Reisolation from *S. arenicola* CNY-014

Pure **1** was obtained from the CBY014-S10 fraction by pTLC purification. The S10 fraction was dissolved in 100 µL of MeOH, applied to a pTLC plate (5 cm × 20 cm), and eluted with a 1:9 mixture of MeOH:EtOAc. Six bands (CBY014-S10A to CBY014-S10F) were obtained. Compound **1** was found in fraction CBY014-S10E. Repetition of the pTLC purification eluting with a 1:10 mixture of MeOH:acetone provided pure **1** (580 µg). This material was used for NMR analyses and structure elucidation studies. Samples of rifamycin B (Sigma Aldrich, SIAL-R0900000) and rifamycin S (TCI America, R02001G) were obtained commercially to assist in the structure elucidation effort.

### Structure elucidation of 1

NMR data were acquired with a Bruker Avance III 600 MHz spectrometer equipped with a 1.7mm cryoprobe. Chemical shifts were referenced using the corresponding solvent signals (*δ*_H_ 7.26 and *δ*_C_ 77.0 for CDCl_3_, δ_H_ 3.31 and δ_C_ 49.0 for CD_3_OD). The NMR spectra were processed using Mnova 11.0 (Mestrelab Research) or TopSpin 3.6 (Bruker Biospin) software.

### Data availability

The authors declare that all data and findings of this study are available within the article and its Supplementary Information file.

## Acknowledgments

M.A. acknowledges financial support from the Scottish Universities Life Sciences Alliance (SULSA) and the Medical Research Council (MRC, J54359) Strategic Grant. M.A, N.T.P, J.A. and R.F., acknowledge financial support from the Wellcome Trust (Grant 201531/Z/16/Z). J.A and F.S acknowledge financial support from BBSRC ISP grants BBS/E/D/20002173 and BBS/E/D/20002174. We thank Scott Franzblau (UIC) and Baoje Wan (UIC) for conducting the Mtb assays.

## Author contributions

J.R.F. and M.A. conceived the assay and screen. N.T.P., J.A., F.A.S., J.R.F. and M.A. conducted the assay. N.T.P., J.A., F.A.S., J.R.F. and M.A. interpreted the assay and identified the hit test fraction. W.F. provided the test fractions. R.C. and W.F. cultured the *S. arenicola* strains and provided extracts for compound isolation. J.J.L. conducted the compound isolation and purification efforts. J.J.L. collected the NMR and MS data. M.S.B., J.J.L., B. M. D. and W.F. evaluated the NMR data and conducted the structural assignments. J.J.L. and M.A. wrote the manuscript. D.A.M. contributed to the biosynthetic discussion. All authors discussed the results and edited the manuscript prior to submission.

## Additional information

Supplementary information. The online version contains supplementary information available online at

## Supplementary Information

**Supplementary Table 1.**
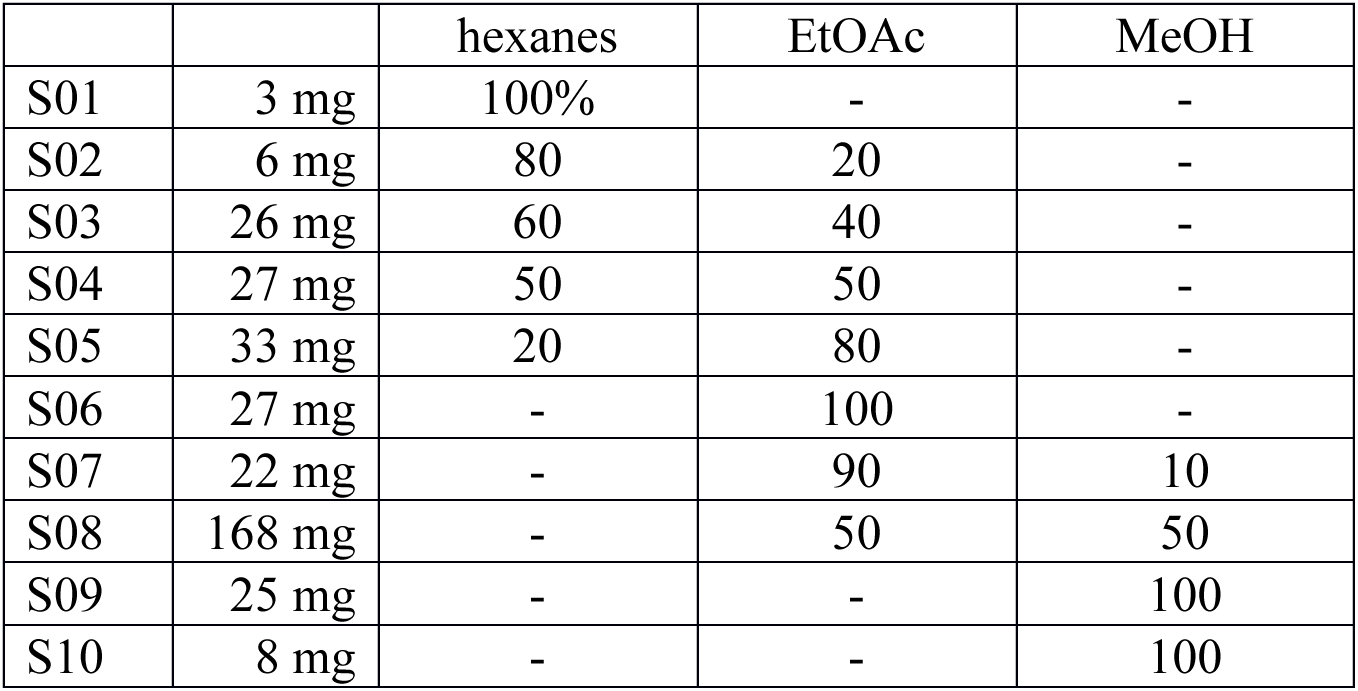
Fractionation method and materials obtained

|  |  | hexanes | EtOAc | MeOH |
| --- | --- | --- | --- | --- |
| S01 | 3 mg | 100% | - | - |
| S02 | 6 mg | 80 | 20 | - |
| S03 | 26 mg | 60 | 40 | - |
| S04 | 27 mg | 50 | 50 | - |
| S05 | 33 mg | 20 | 80 | - |
| S06 | 27 mg | - | 100 | - |
| S07 | 22 mg | - | 90 | 10 |
| S08 | 168 mg | - | 50 | 50 |
| S09 | 25 mg | - | - | 100 |
| S10 | 8 mg | - | - | 100 |

**Supplementary Table 2.** NMR data for rifamycin analogue **1** in CD_3_OD

| | $\delta_{\text{H}}$, mult ($J$ in Hz) | $\delta_{\text{C}}$<sup>1,2</sup> | <sup>1</sup>H, <sup>1</sup>H-CCOSY | <sup>1</sup>H, <sup>1</sup>H-NOESY<sup>3</sup> | <sup>1</sup>H, <sup>13</sup>C-HMBC |
| --- | --- | --- | --- | --- | --- |
| 1 |  | ND<sup>4</sup> |  |  |  |
| 2 |  | ND<sup>4</sup> |  |  |  |
| 3 |  | 146.6<sup>6</sup> |  |  |  |
| 4 |  | 188.9<sup>6</sup> |  |  |  |
| 5 |  | ND<sup>4</sup> |  |  |  |
| 6 |  | 172.0 |  |  |  |
| 7 |  | 108.6 |  |  |  |
| 8 |  | 173.5 |  |  |  |
| 9 |  | ND<sup>4</sup> |  |  |  |
| 10 |  | ND<sup>4</sup> |  |  |  |
| 11 |  | 198.4 |  |  |  |
| 12 |  | 109.1 |  |  |  |
| 13 | 1.67, s | 21.7 | – | – | 11,12 |
| 14 | 2.10, s | 6.8 | – | – | 6,7,8 |
| 15 |  | 171.7 |  |  |  |
| 16 |  | 124.8 |  |  |  |
| 17 | 6.21, d (11.5) | 133.6 | 18,30 | 19,30 | 15w,19,30 |
| 18 | 7.11, dd (16.0, 10.7) | 128.5 | 17,19 | 21,31 | 17,20w |
| 19 | 6.07, dd (16.2, 7.4) | 139.6 | 18,20 | 17,31w | 17,20w |
| 20 | 2.34, dt (9.9, 7.0) | 38.6 | 19w,21,31 | 32 | 19w |
| 21 | 4.05, d (11.2) | 75.6 | 20,22 | 18,24,31,33 | 32 |
| 22 | 1.91, m | 34.2 | 21w,23w,32 | 31,33 | – |
| 23 | 3.46, dd (10.3, 2.5) | 78.4 | 22,24 | 25w,32,33 | 21w |
| 24 | 1.56, m | 39.8 | 23,33 | 21,34 | – |
| 25 | 3.71, d (10.7) | 72.2 | 26 | 23,27w,33w,34 | 23w,26w,33 |
| 26 | 1.33, m | 42.2 | 25,34 | 33 | – |
| 27 | 4.42, d (7.0) | 68.3 | 26,28,29w | 25,29,34w | 26,29,34 |
| 28 | 5.31, dd (12.7, 7.0) | 124.9 | 27,29 | 26,34 | 29 |
| 29 | 6.24, dd (12.7, 1.4) | 141.9 | 27w,28 | 26,27,34 | 27w, 28 |
| 30 | 2.01, s | 20.5 | – | – | 15,17 |
| 31 | 0.98, d (6.7) | 18.3 | 20 | 21,22 | 19,20,21 |
| 32 | 1.05, d (6.9) | 10.8 | 22 | 20,23 | 21,22,23 |
| 33 | 0.74, d (6.9) | 9.1 | 24 | 21,22,23,26 | 23,24,25 |
| 34 | -0.05, d (6.7) | 9.9 | 26 | 24,25,28w | 25,26,27 |
| 35 | 3.04, m | 56.5 |  | 37 | 3,4,36 |
| 36 |  | 208.8 |  |  |  |
| 37 | 2.29, s | 32.9 |  | 35 | 36 |
| NH | 8.54, s<sup>5</sup> |  |  |  |  |

<sup>1,2</sup> Assignments obtained <sup>1</sup> from <sup>1</sup>H, <sup>13</sup>C-HSQC or <sup>2</sup> from <sup>1</sup>H, <sup>13</sup>C-HMBC spectra.
<sup>3</sup> NOEs from protons on neighbouring carbons are not tabulated.
<sup>4</sup> Not determined. Sample sizes were too small to collect directly detected <sup>13</sup>C NMR spectra, nor were we able to obtain sufficient sensitivity to assign these carbons from <sup>1</sup>H, <sup>13</sup>C-HMBC spectra.
<sup>5</sup> This assignment was tentative and could arise from an impurity such as formate
<sup>6</sup> These assignments could be swapped.

**Supplementary Table 3.** NMR data for rifamycin B in CD_3_OD

| | $\delta_{\text{H}}$, mult (*J* in Hz) | $\delta_{\text{C}}$ | $^1\text{H}$, $^1\text{H}$-COSY | $^1\text{H}$, $^{13}\text{C}$-HMBC |
| --- | --- | --- | --- | --- |
| 1 |  | 147.9<sup>2</sup> |  |  |
| 2 |  | 123.4<sup>2</sup> |  |  |
| 3 | 7.37, bs | 109.1 |  | – |
| 4 |  | 147.9<sup>1</sup> |  |  |
| 5 |  | 103.5<sup>2</sup> |  |  |
| 6 |  | 159.7<sup>2</sup> |  |  |
| 7 |  | 107.6<sup>2</sup> |  |  |
| 8 |  | 119.0<sup>2</sup> |  |  |
| 9 |  | 107.6<sup>2</sup> |  |  |
| 10 |  | 115.3<sup>2</sup> |  |  |
| 11 |  | 175.4<sup>1</sup> |  |  |
| 12 |  | 110.3<sup>1</sup> |  |  |
| 13 | 1.76, s | 22.2 |  | 12 |
| 14 | 2.19, s | 7.6 |  | 7,9,11 |
| 15 |  | 172.7<sup>1</sup> |  |  |
| 16 |  | 132.1<sup>1</sup> |  |  |
| 17 | 6.31, d (10.5) | 134.2 | 18,30w | 15,18w,19w,30 |
| 18 | 6.43, dd (15.6, 10.9) | 125.3 | 17,19 | 17w,20 |
| 19 | 6.01, dd (15.5, 6.0) | 141.7 | 18,20 | 17,20w,21 |
| 20 | 2.34, m | 39.6 | 19w,21,31 | 18w,19w,21w,31w |
| 21 | 3.86, d (8.8) | 73.6 | 20,22w | 19w,20w,32 |
| 22 | 1.78, m | 34.4 | 32 | 23w,32 |
| 23 | 3.09, dd (10.2, 2.2) | 78.1 | 22w,24 | 21,22w,24 |
| 24 | 1.45, dq (13.9, 7.2) | 39.1 | 23,33 | 23.33 |
| 25 | 5.12, d (10.7) | 74.9 | 26 | 23w,24w,26,33,34,35 |
| 26 | 1.20, m | 41.0 | 25,34 | 25w |
| 27 | 3.43, d (7.0) | 78.5 | 28 | 25,26,28,29,34,37 |
| 28 | 5.08, dd (12.7, 7.0) | 119.6 | 27,29 | 27w,29 |
| 29 | 6.20, dd (12.7, 1.0) | 144.0 | 28 | 12,27,28 |
| 30 | 2.09, s | 20.4 | 17w | 15,16,17,18w,19w |
| 31 | 0.92, d (7.0) | 17.9 | 20 | 20,21 |
| 32 | 0.99, d (7.0) | 11.4 | 22 | 21,22,23 |
| 33 | 0.60, d (6.9) | 9.4 | 24 | 23,24,25 |
| 34 | -0.34, d (6.9) | 9.4 | 26 | 25.26.27 |
| 35 |  | 172.6<sup>1</sup> |  |  |
| 36 | 2.02, s | 20.8 |  | 35 |
| 37 | 3.03, s | 57.1 | – | 27 |
| 38 | 4.74, s | 67.7 |  | 4,39 |
| 39 |  | 171.9<sup>1</sup> |  |  |
| NH | 7.92, d (0.6) |  |  |  |

<sup>1</sup> Assignments obtained from $^1\text{H}$ , $^{13}\text{C}$ -HMBC spectrum<sup>2</sup> Assignments based on chemical shift predictions and literature precedent

**Supplementary Table 4.** NMR data for rifamycin S in CD_3_OD^3^

| | $\delta_{\text{H}}$, mult (*J* in Hz) | $\delta_{\text{C}}$ | $^1\text{H}$, $^1\text{H}$-COSY | $^1\text{H}$, $^{13}\text{C}$-HMBC |
| --- | --- | --- | --- | --- |
| 1 |  | 184.0<sup>1,2</sup> |  |  |
| 2 |  | 141.2<sup>2</sup> |  |  |
| 3 | 7.63, s | 118.3 | – | 2,4,5,10 |
| 4 |  | 185.9<sup>1,2</sup> |  |  |
| 5 |  | 112.2<sup>1,2</sup> |  |  |
| 6 |  | 168.0<sup>1,2</sup> |  |  |
| 7 |  | 117.0<sup>1,2</sup> |  |  |
| 8 |  | 173.9<sup>1,2</sup> |  |  |
| 9 |  | 112.0<sup>1,2</sup> |  |  |
| 10 |  | 131.9<sup>1,2</sup> |  |  |
| 11 |  | 193.9<sup>1</sup> |  |  |
| 12 |  | 109.5<sup>1</sup> |  |  |
| 13 | 1.70, s | 22.2 | – | 11,12,29w |
| 14 | 2.31, s | 7.7 | – | 1w,5,6,7,8,9,10 |
| 15 |  | 172.0<sup>1</sup> |  |  |
| 16 |  | 132.3<sup>1</sup> |  |  |
| 17 | 6.30, d (10.5) | 134.3 | 18,30 | 15,19,30 |
| 18 | 6.26, m | 125.8 | 17,19 | 16,17,20 |
| 19 | 5.91, dd (14.3, 6.5) | 142.2 | 18,20 | 15w,17,20,21,30w,31 |
| 20 | 2.33, m | 40.1 | 19w,21,31 | 18,19,21,31 |
| 21 | 3.69, dd (9.8, 1.9) | 74.5 | 20,22w | 19,20,21w,22w,23,32 |
| 22 | 1.78, m | 34.0 | 21w,32 | 20w,23,32 |
| 23 | 3.10, d (2.1) | 77.9 | 22w,24 | 21,22,24,32 |
| 24 | 1.51, m | 38.7 | 23,33 | 22w,23,33 |
| 25 | 4.93, dd (10.5, 1.7) | 73.9 | 24w,26 | 23,24,26,27,33,34w,35 |
| 26 | 1.83, dtd (10.4, 7.1, | 38.8 | 25,27w,34 | 25,27w,28w,34 |
| 27 | 3.38, dd (7.9, 3.1) | 82.3 | 26w,28 | 25,26,28,29,34,37 |
| 28 | 5.92, dd (12.6, 7.9) | 118.1 | 27,29 | 12w,26w,27w,29 |
| 29 | 6.20, dd (12.6, 0.8) | 145.6 | 28 | 12,27,28 |
| 30 | 2.04, s | 20.2 | 17 | 15,16,17,18w,19w,31w |
| 31 | 0.86, d (6.9) | 17.5 | 20 | 20,21 |
| 32 | 0.99, d (7.0) | 11.4 | 22 | 21,22,23 |
| 33 | 0.69, d (6.9) | 9.3 | 24 | 23,24,25 |
| 34 | 0.09, d (7.1) | 12.0 | 26 | 25,26,27 |
| 35 |  | 172.9<sup>1</sup> |  |  |
| 36 | 1.95, s | 21.1 | – | 25w,35 |
| 37 | 3.08, s | 56.9 | – | 27 |

<sup>1</sup> Assignments obtained from $^1\text{H}$ , $^{13}\text{C}$ -HMBC spectrum<sup>2</sup> Assignments based on chemical shift predictions and literature precedent<sup>3</sup> Contains impurity with 1.15, d (6.1), 25.3 and 3.93, h (6.2). 64.7

**Supplementary Table 5.**
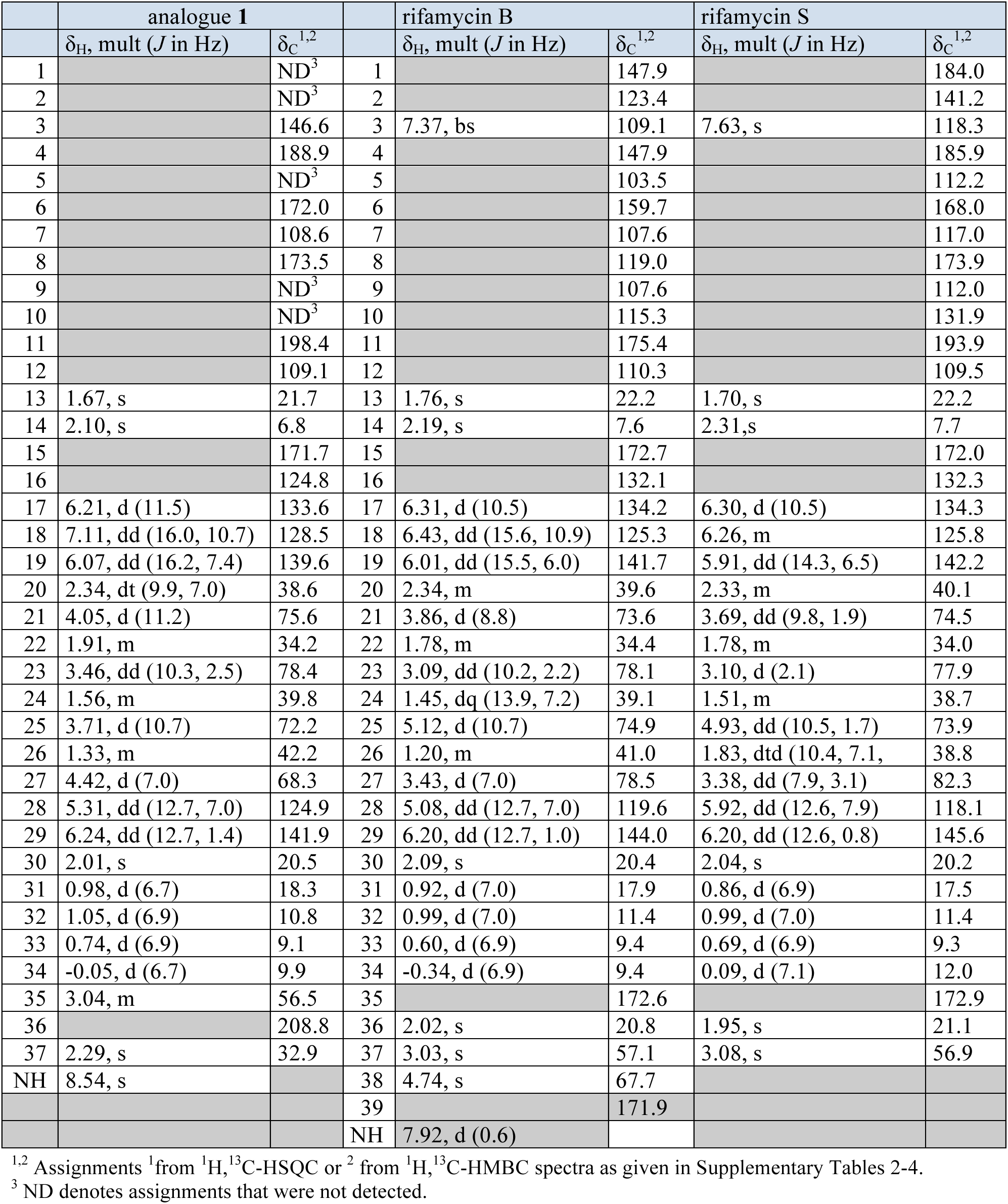
Aligned ^1^H and ^13^C NMR data for **1**, rifamycin B, and rifamycin S in CD_3_OD

**Supplementary Table 6.** Parameters used for image analysis using Acapella

| **General Parameters** | **Value** |
| --- | --- |
| Channel number for nuclear channel | 4 |
| Channel number for cytoplasm channel | 3 |
| Channel number for spot channel | 1 |
| Spot detection area | WholeCell |
| Remove Border Objects | Yes |
| Minimum roundness of cells (1 = perfect circle) | 0.8 |
| **Nuclei Detection** |  |
| Nuclei Detection Algorithm | H |
| Threshold Adjustment | 1 |
| Minimum Nuclei Distance | 10 |
| Nuclear Splitting Adjustment | 8 |
| Individual Threshold Adjustment | 0.45 |
| Minimum Nuclear Area | 100 |
| Minimum Nuclear Contrast | 0.3 |
| **Nucleus Region** |  |
| NucleusOuterBorderShift | 0 |
| NucleusInnerBorderShift | unlimited |
| **Cytoplasm Detection** |  |
| Cytoplasm Threshold Adjustment | 0.7 |
| Cytoplasm Individual Threshold Adjustment | 0.5 |
| **Cytoplasm Region** |  |
| CytoplasmOuterBorderShift | 0 |
| CytoplasmInnerBorderShift | 0 |
| NucleusBorderShift | unlimited |
| **Membrane Region** |  |
| CellOuterBorderShift | -1 |
| CellInnerBorderShift | 1 |
| **Object Region** |  |
| ObjectBorderShift | 2 |
| **Spot Detection** |  |
| SpotMinimumDistance | 2 |
| SpotPeakRadius | 0 |
| SpotReferenceRadius | 3 |
| SpotMinimumContrast | 0.2 |
| SpotMinimumToCellIntensity | 0.7 |

**Supplementary Fig. 1.**
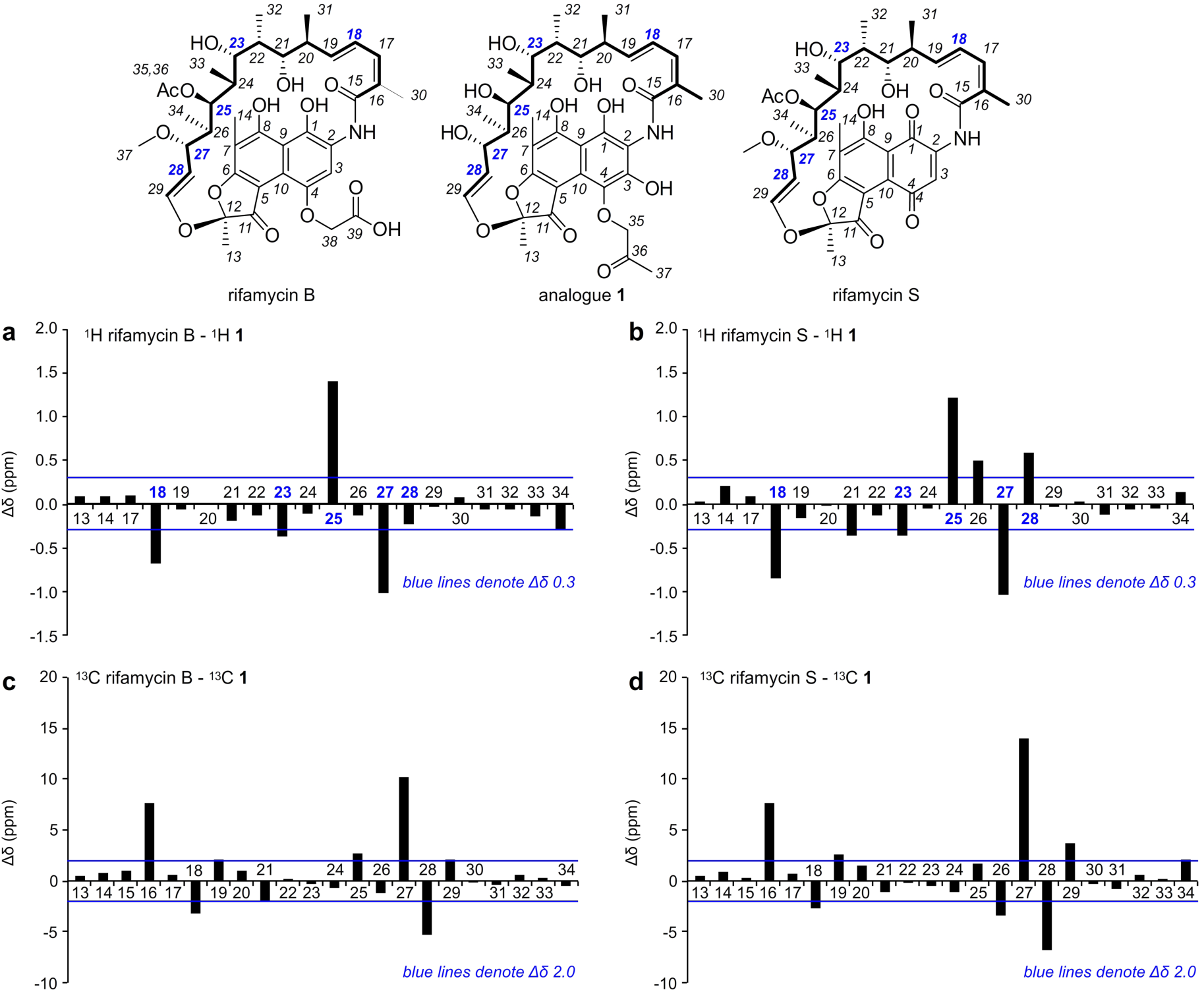
NMR peak shift analyses. Comparison of the ^1^H (top row) and ^13^C NMR (bottom row) peaks of **1** to rifamycin B (left) and rifamycin S (right column). Blue line denotes Δδ = ±0.3 ppm for ^1^H and Δδ = ±2.0 ppm for ^13^C peaks.

**Supplementary Fig. 2.**
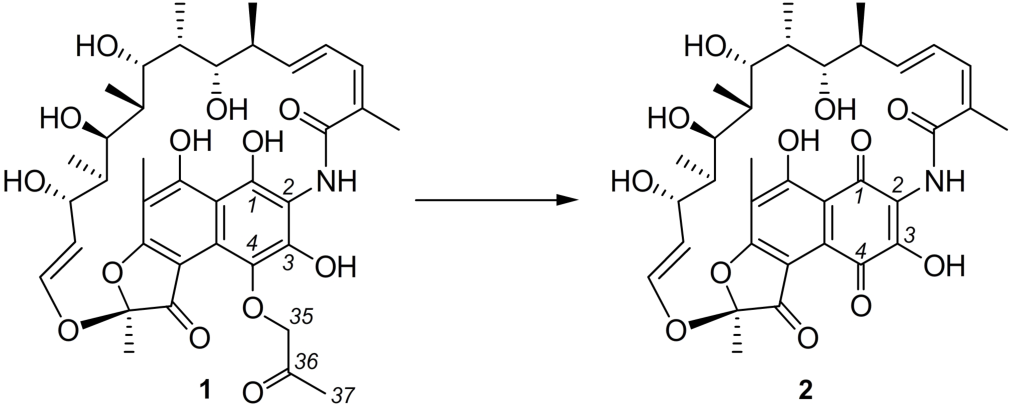
Proposed degradation of shunt metabolite 1. Compound 1 was observed to be very reactive. In MeOH solvent it underwent oxidation followed by methanolysis to form 2. This process is similar to the conversion of rifamycin B to rifamycin S (active form). Traces of 2 were always detected when isolating 1 and could not be removed experimentally under the conditions reported herein. Based on prior SAR data, we would anticipate that the activity of this material arises from 2.

**Supplementary Fig. 3.**
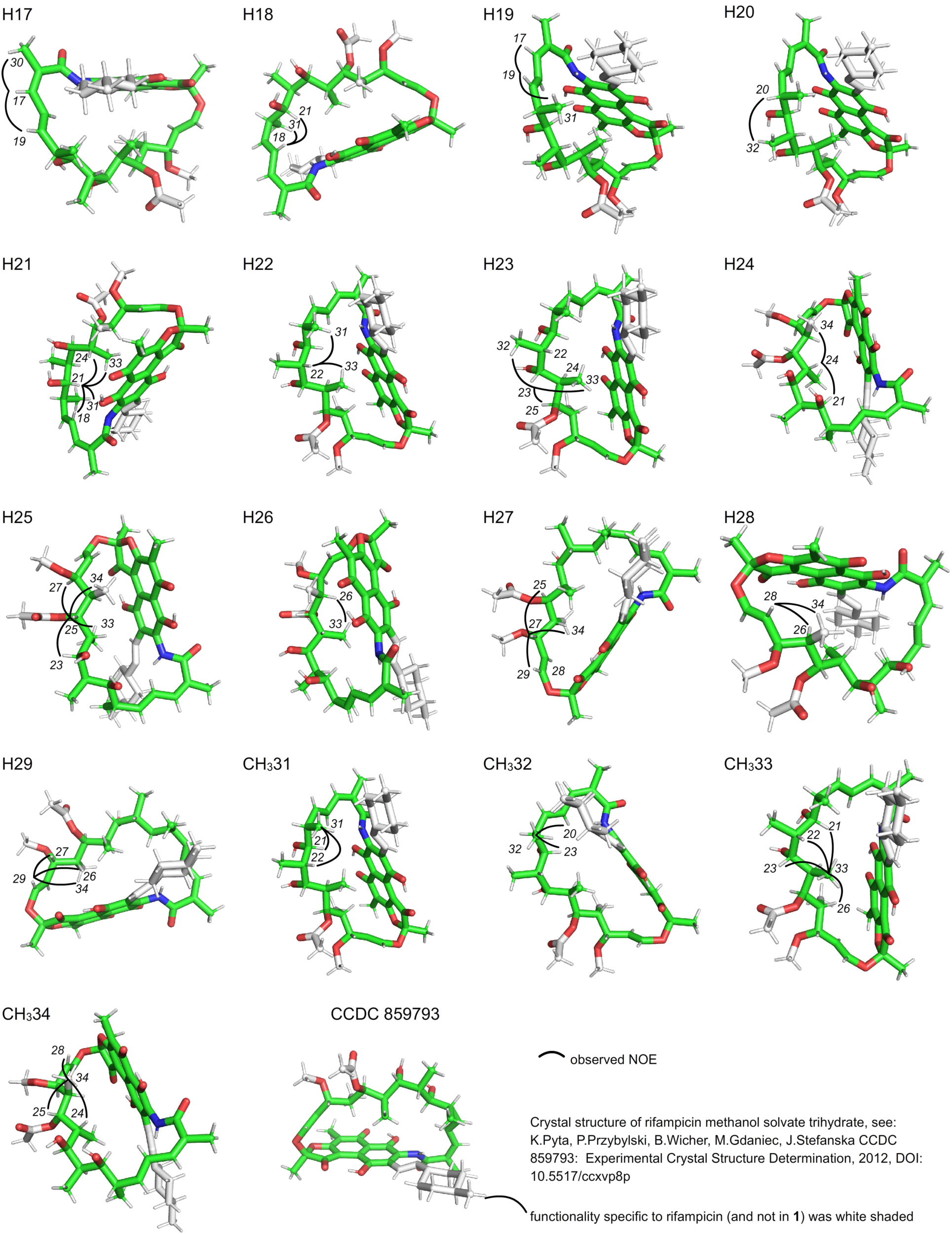
^1^H,^1^H-NOESY correlations observed for 1 shown on the X-ray crystal structure of rifampicin, indicating the stereochemistry is the same.

**Supplementary Fig. 4.**
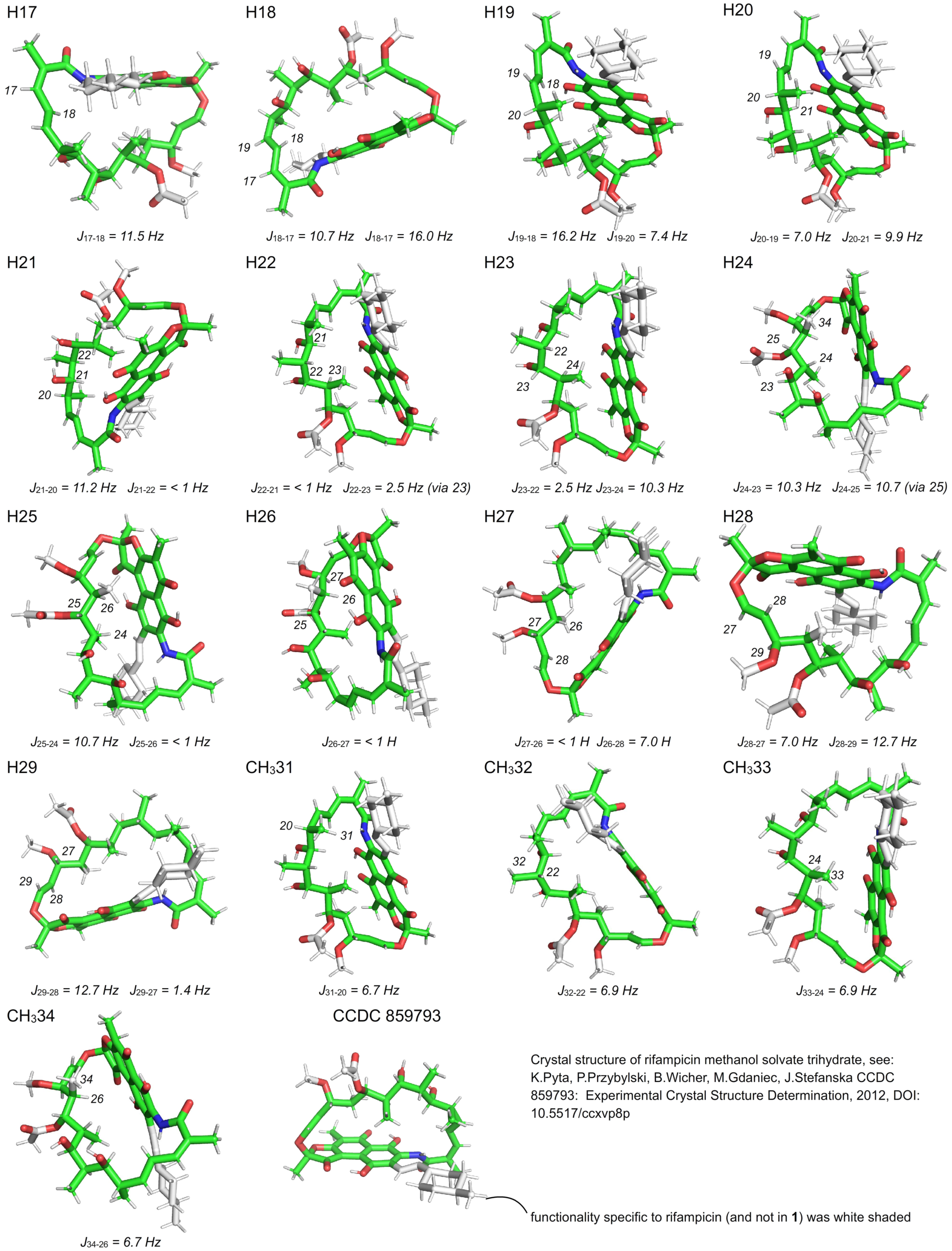
Coupling constants observed for 1 with the X-ray crystal structure of rifampicin oriented to show that the couplings are consistent with the structure.

**Supplementary Fig. 5.**
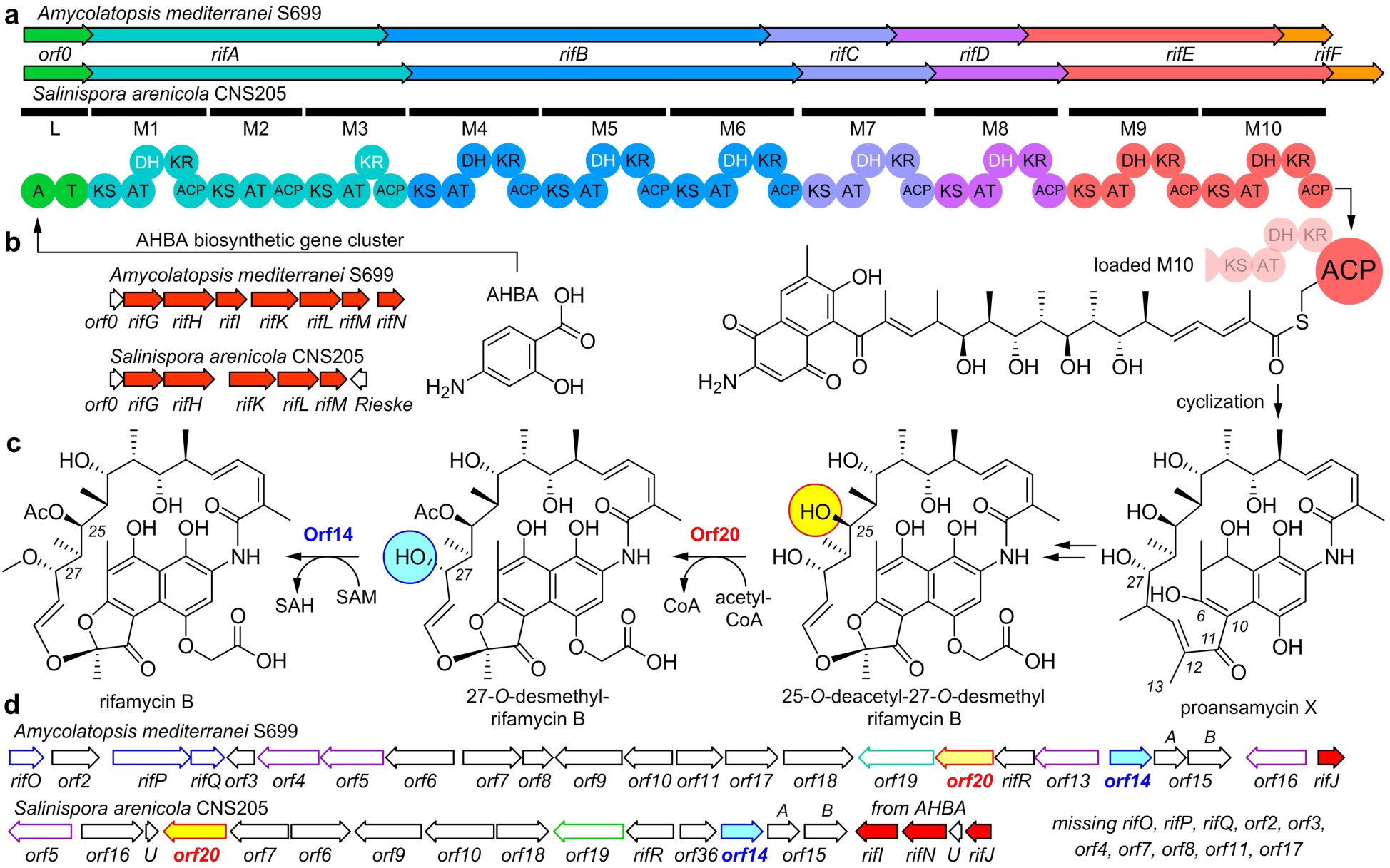
Comparison of the rifamycin BGCs from *A. mediterranei and S. arenicola.* a) The carbon backbone of the rifamycin family is assembled through a polyketide synthase (PKS). PKS assembly involves *orf0* and *rifA*-*rifF* with modular assemblies comprised of AT (acyltransferase), ACP (acyl carrier protein), DH (dehydratase), and KR (ketoreductase) domains. PKS assembly ends at *rifE* where the substrate is released from the terminal ACP in module 10 through amide-forming macrocyclization to deliver proansamysin X. Both strains share very similar PKSs. b) The PKS pathway begins with a 3-amino-5-hydroxy benzoic acid (AHBA) starting unit. In *A. mediterranei*, 8 genes (*orf0*- *rifN*) are located within one cluster, while *rifJ* is located within a region that contains the post-PKS tailoring enzymes. In *S. arenicola*, *rifI* and *rifN* appear proximal to *rifJ*. Amino acid similarities of ∼75% are observed in the PKS and AHBA genes between *A. mediterranei* and *S. arenicola*. c) Conversion to rifamycin SV arises through a multistep process that begins with elaboration of the 5-membered ring between C6 and C12 to form 25-*O*-deacetyl-27-*O*-desmethyl rifamycin B, which undergoes a two-step acetylation and methylation sequence to achieve rifamycin B through CoA-guided acylation by orf20 and SAM-dependent methylation by orf14. d) Noticeably different post-PKS rifamycin modification enzymes are observed in *A. mediterranei* and *S. arenicola,* as shown by this gene comparison.

**Supplementary Fig. 6.**
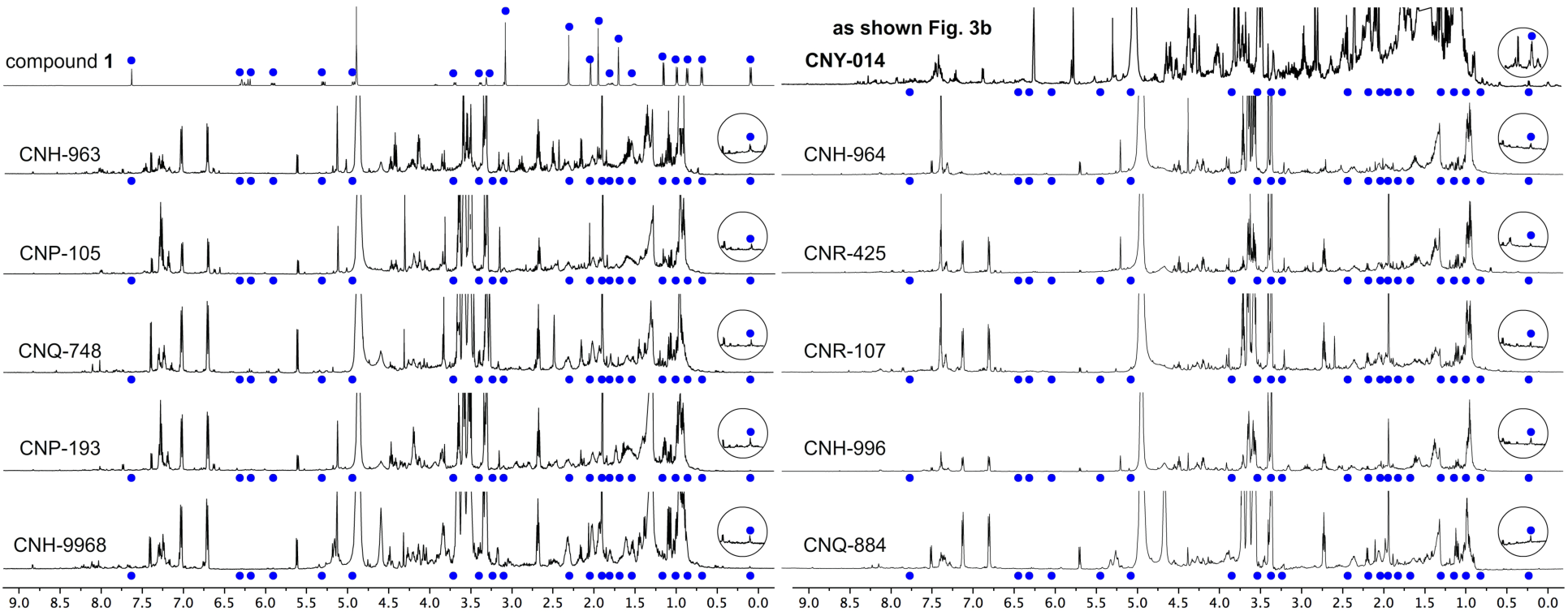
^1^H-NMR spectrum of compound 1 (top row, peaks marked with blue dots) as compared to the crude extracts of ten additional strains of *Salinispora arenicola*. All the extracts show the presence of compound 1.

**Supplementary Fig. 7.**
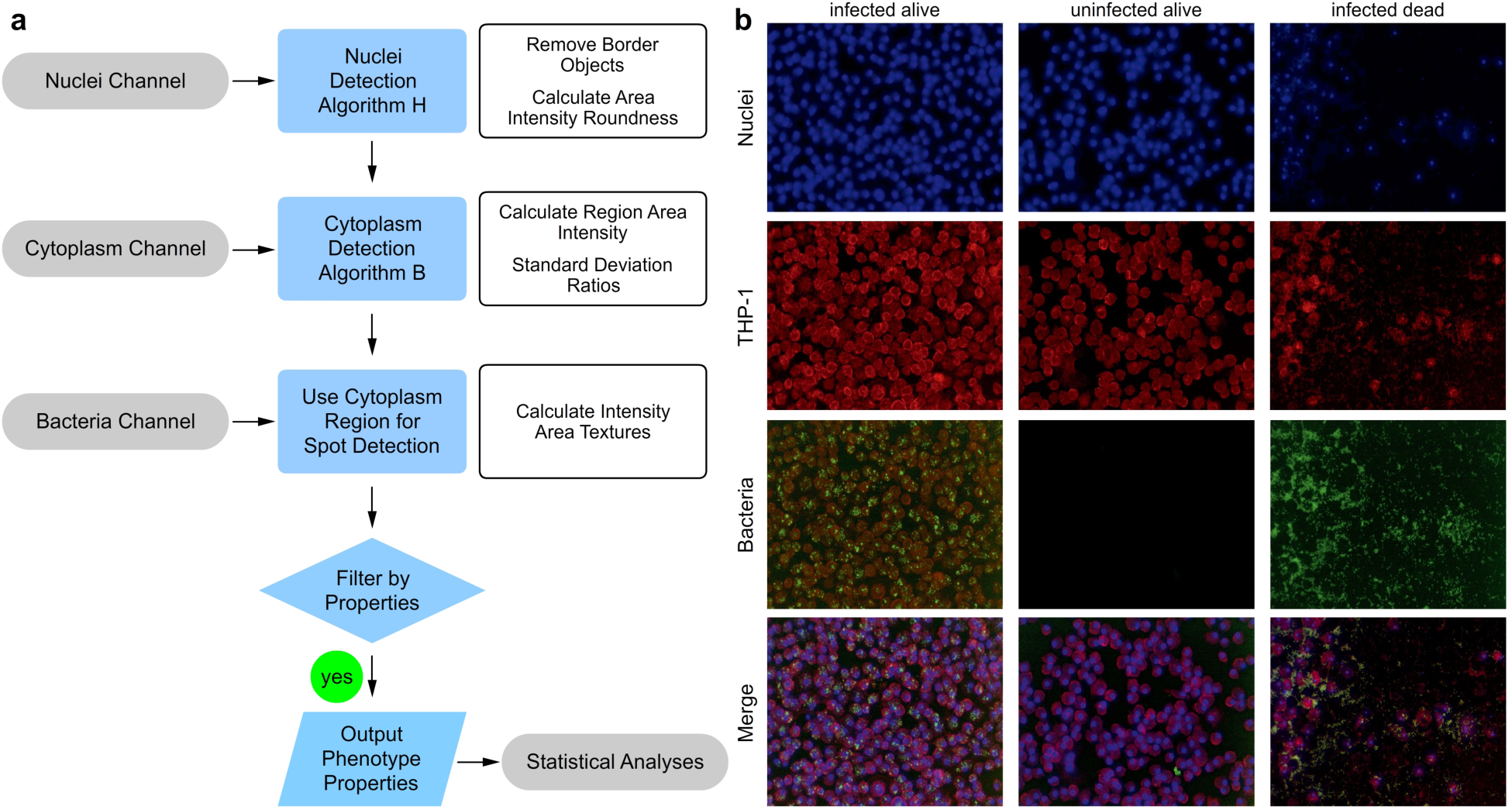
Acapella image analysis. a) Flowchart describing sequential steps used in image analysis. b) Example images visualising *S.aureus* infected THP-1 cells with positive control (left column) and negative control (right). For comparison, uninfected THP-1 cells in media only are shown in the middle column.

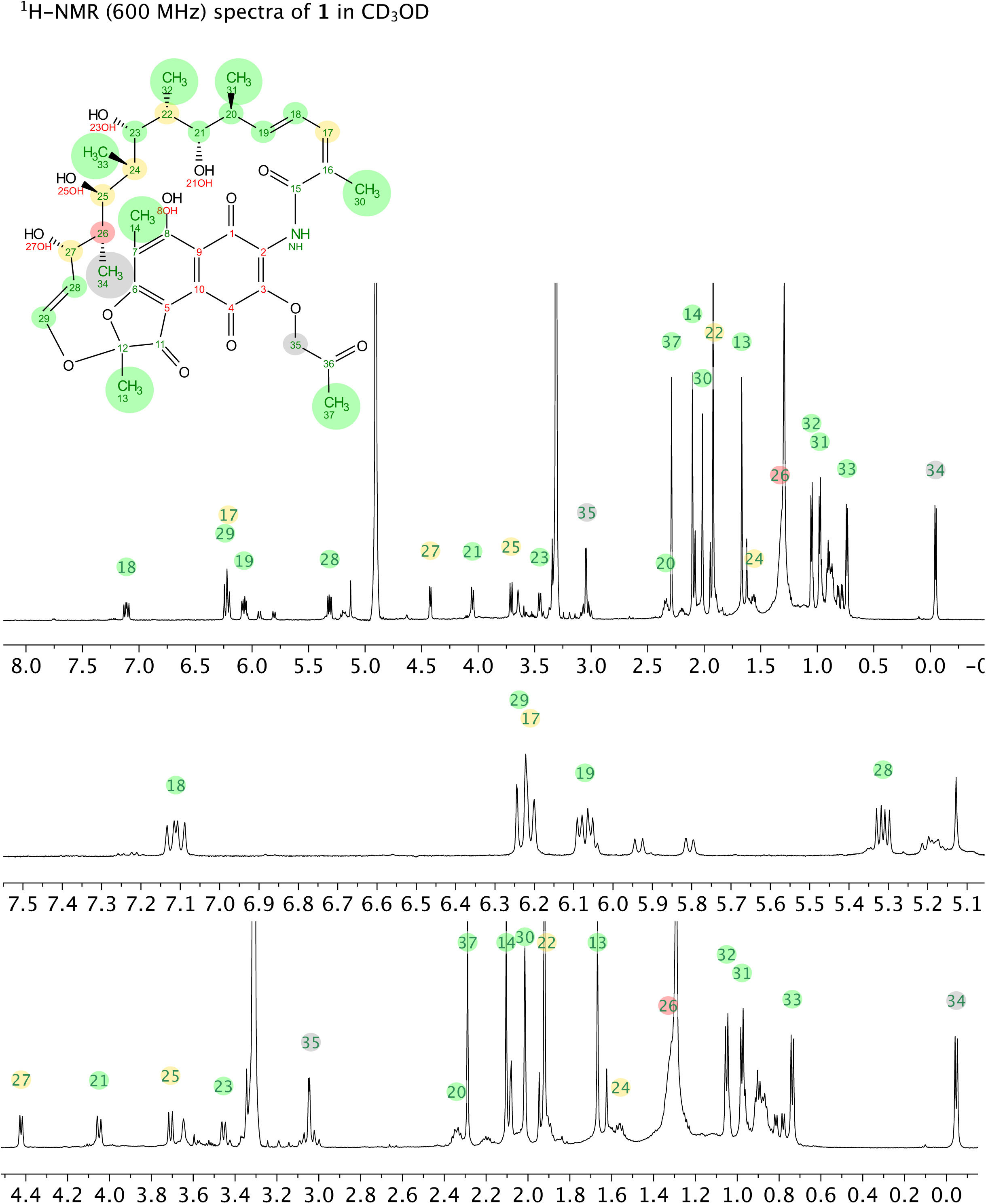
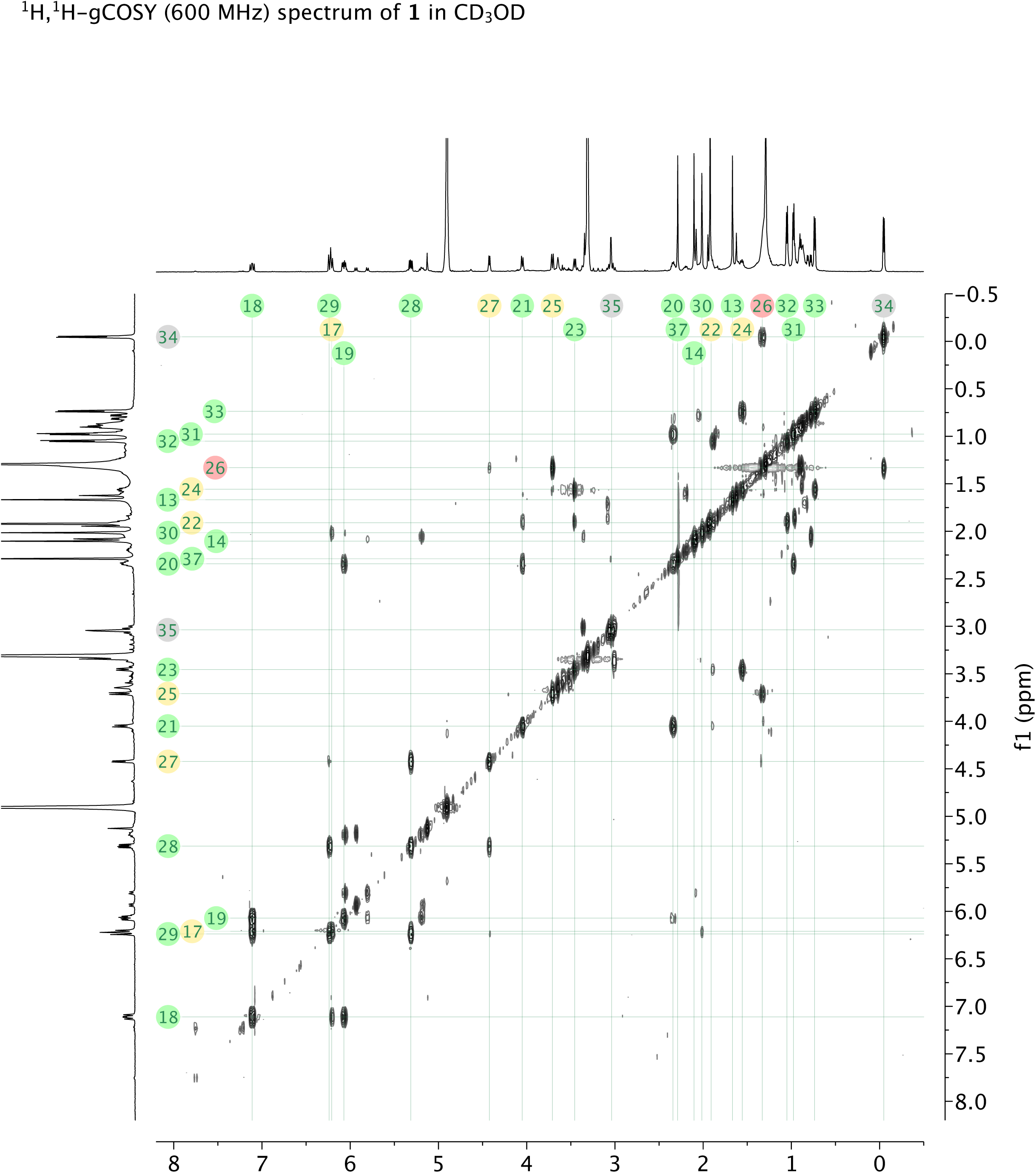
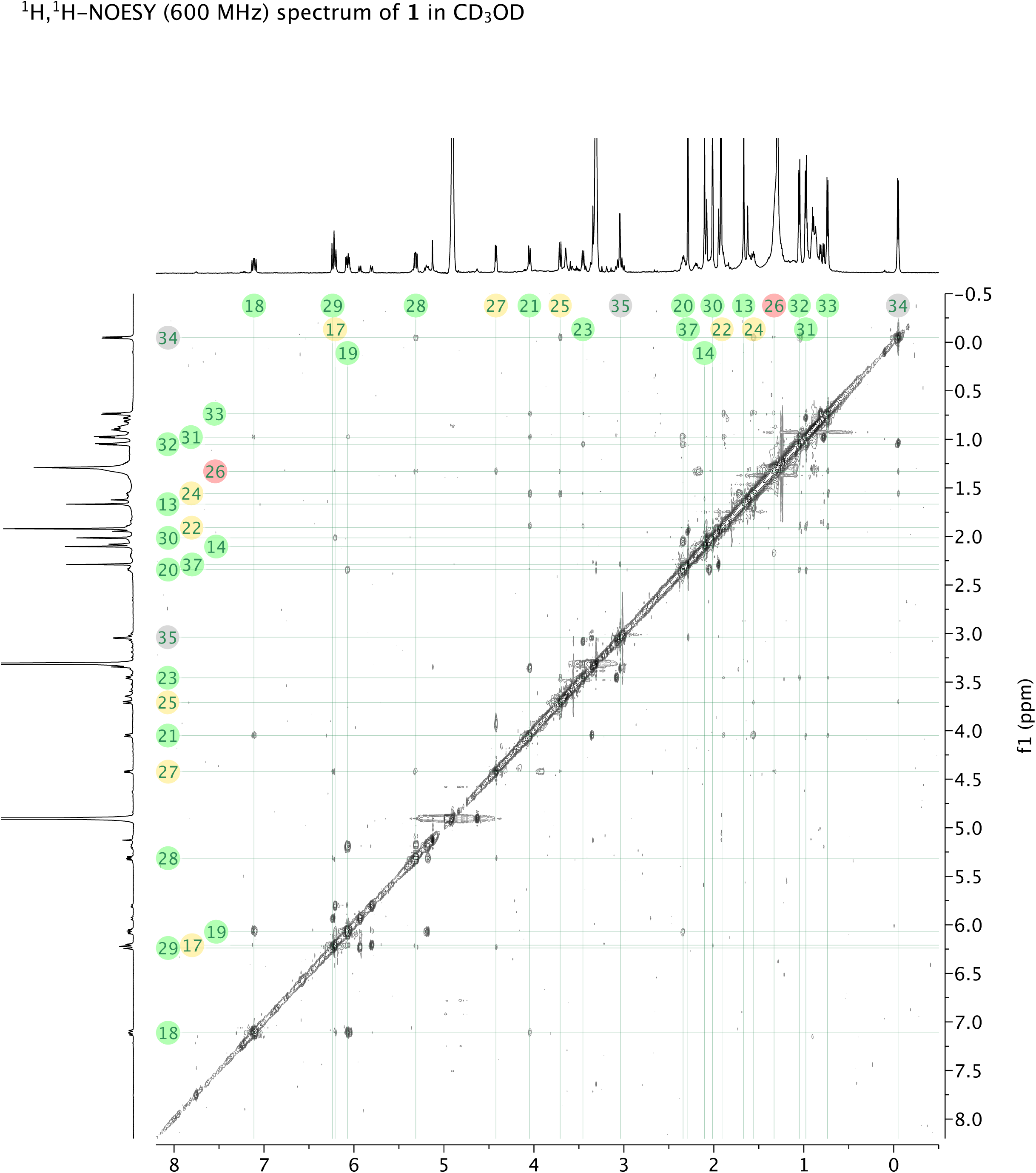
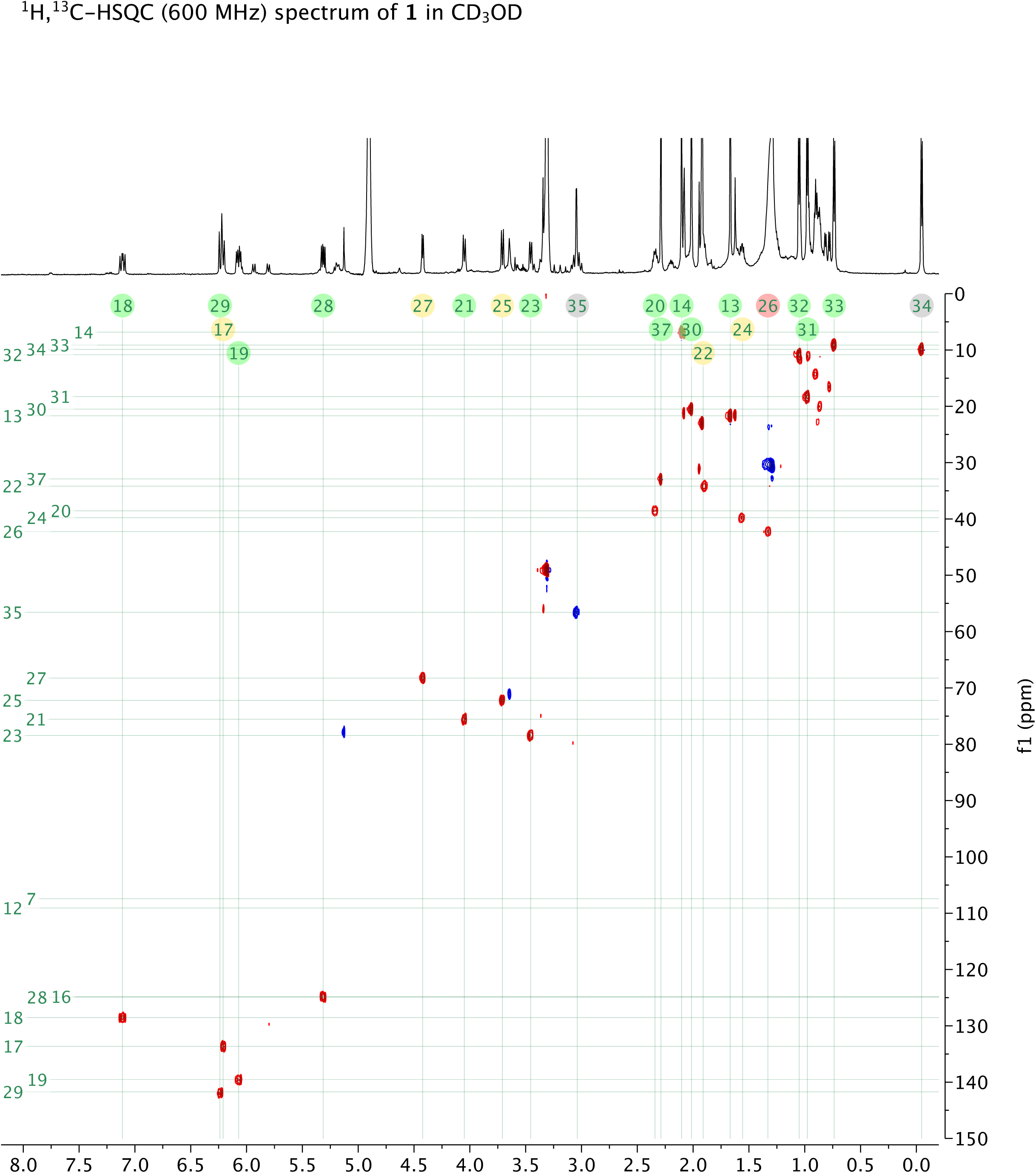
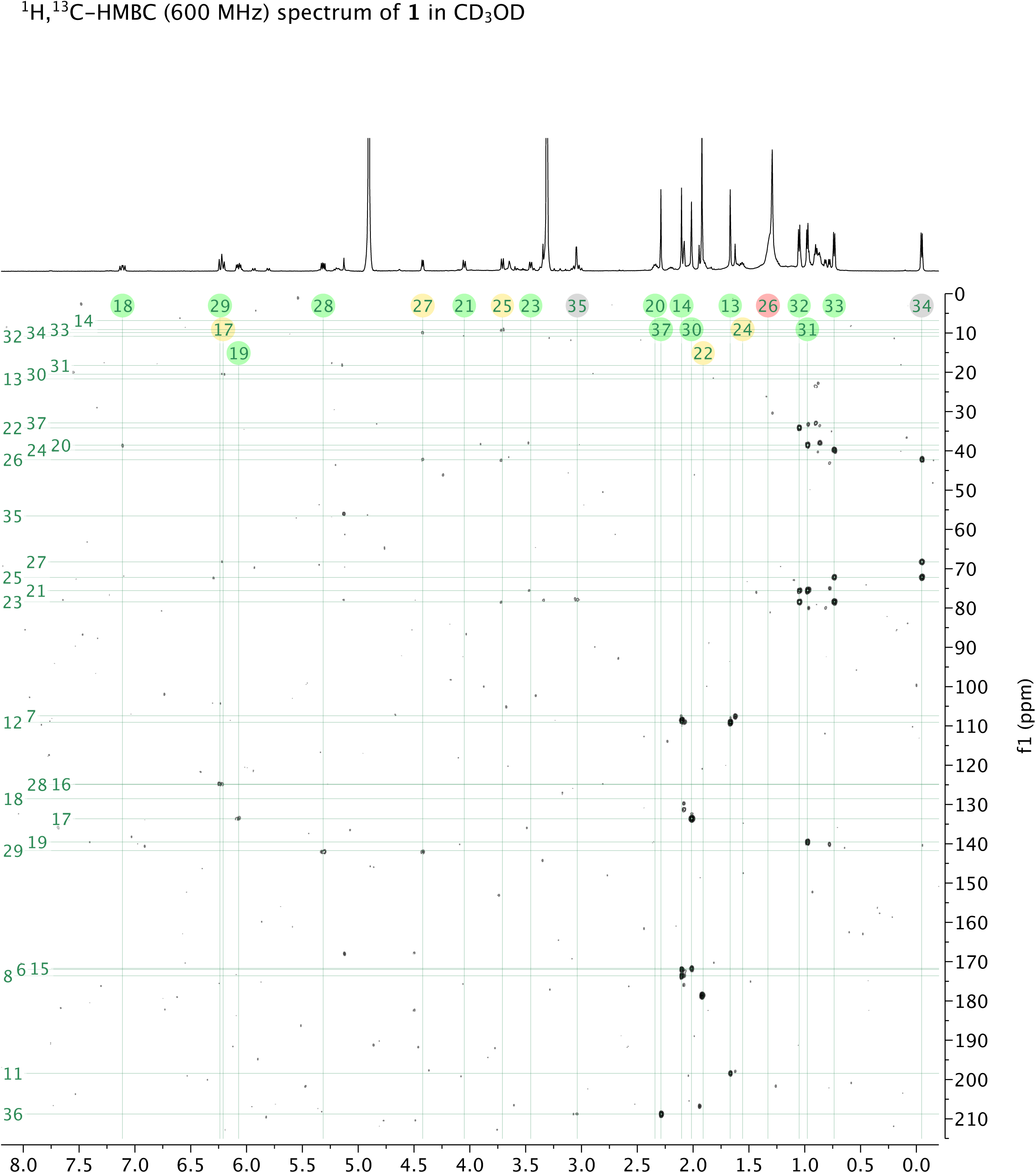
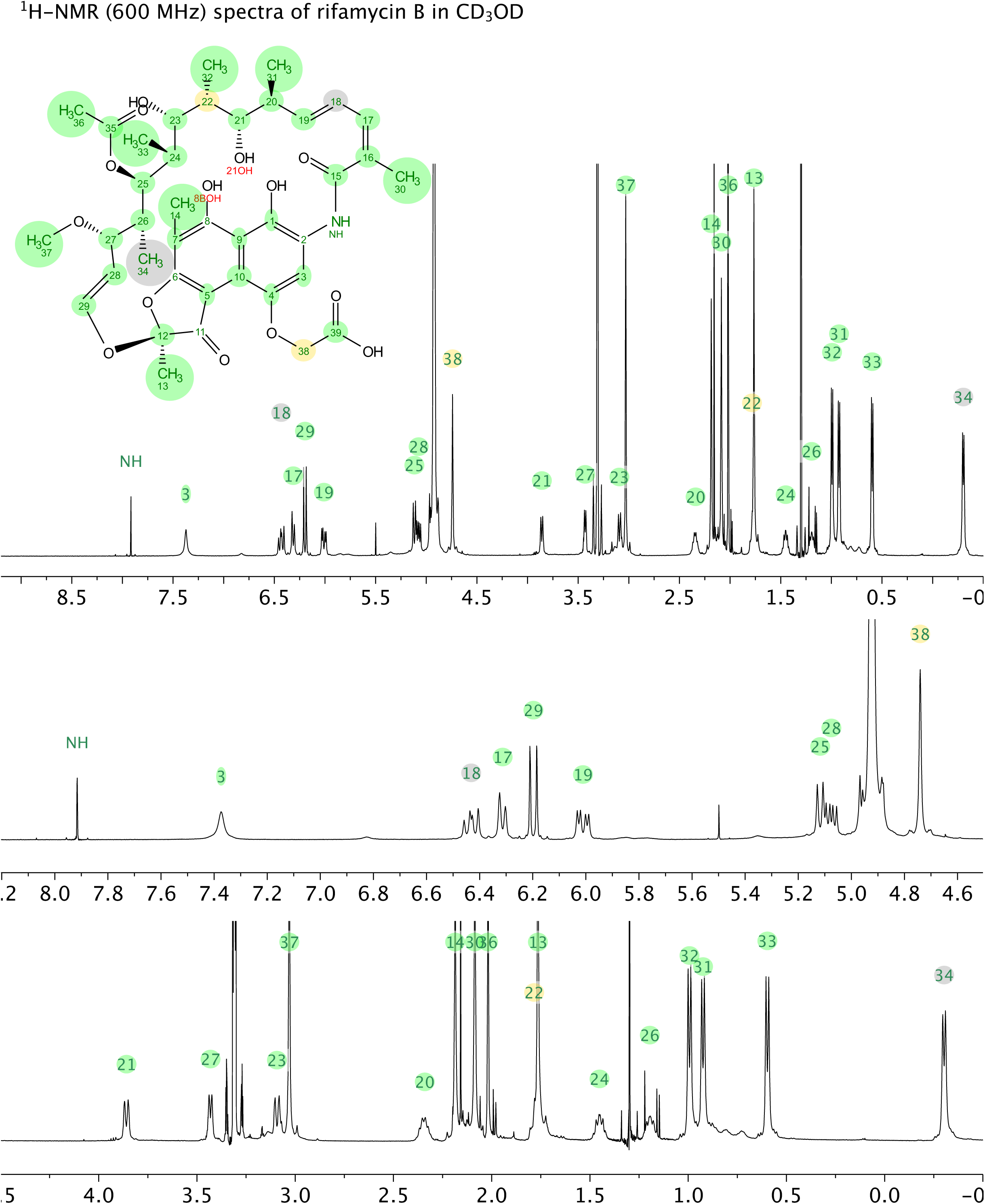
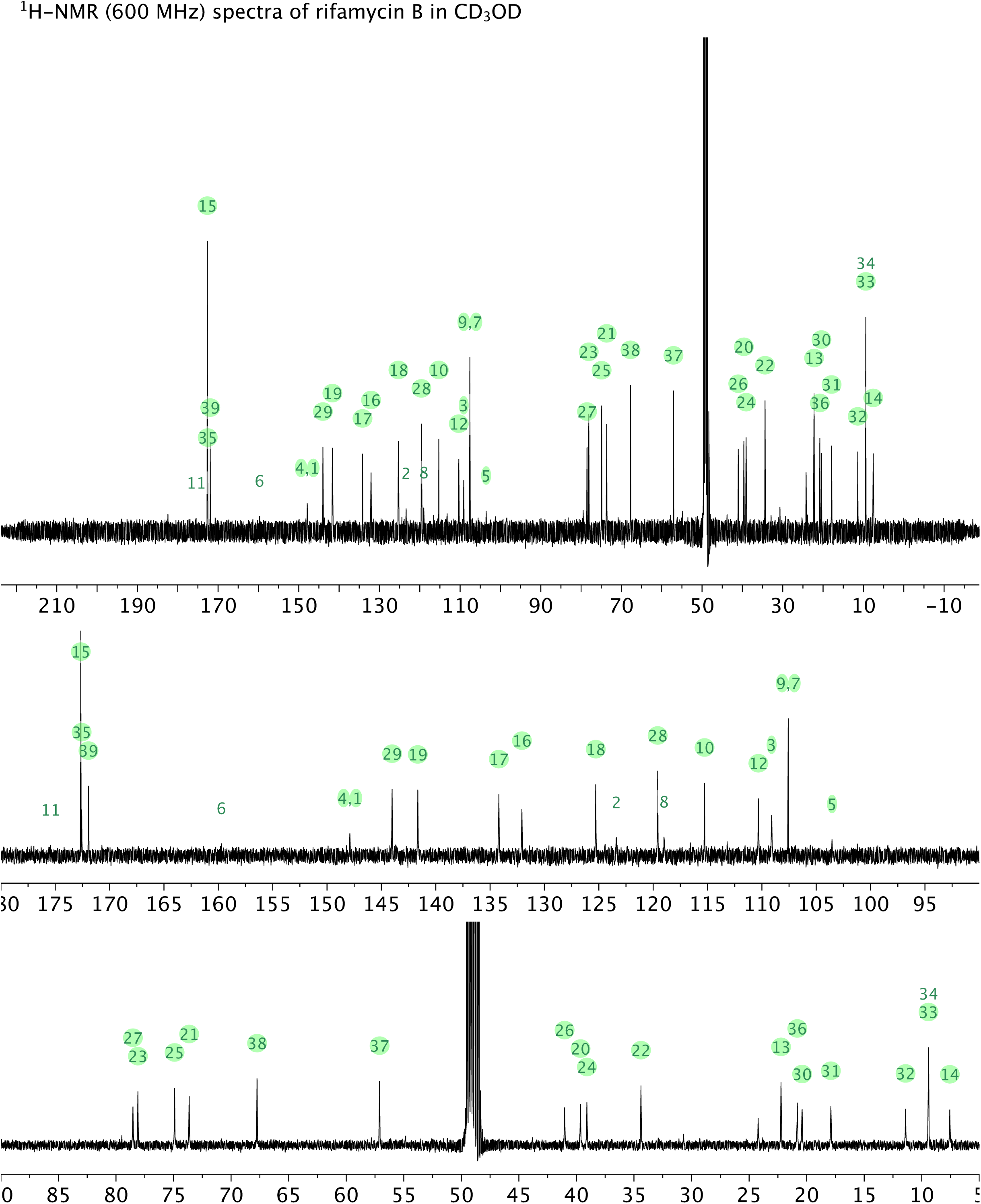
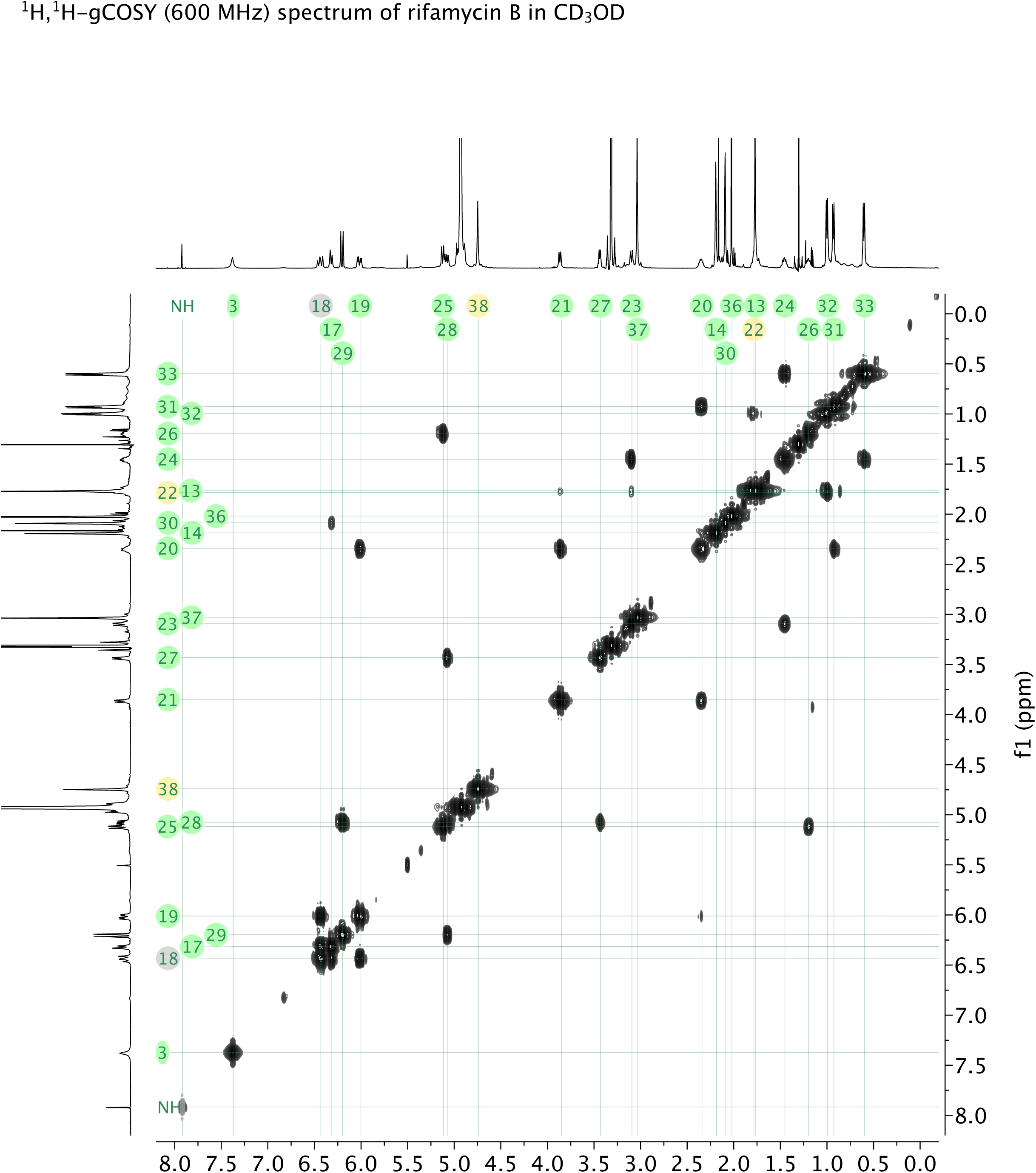
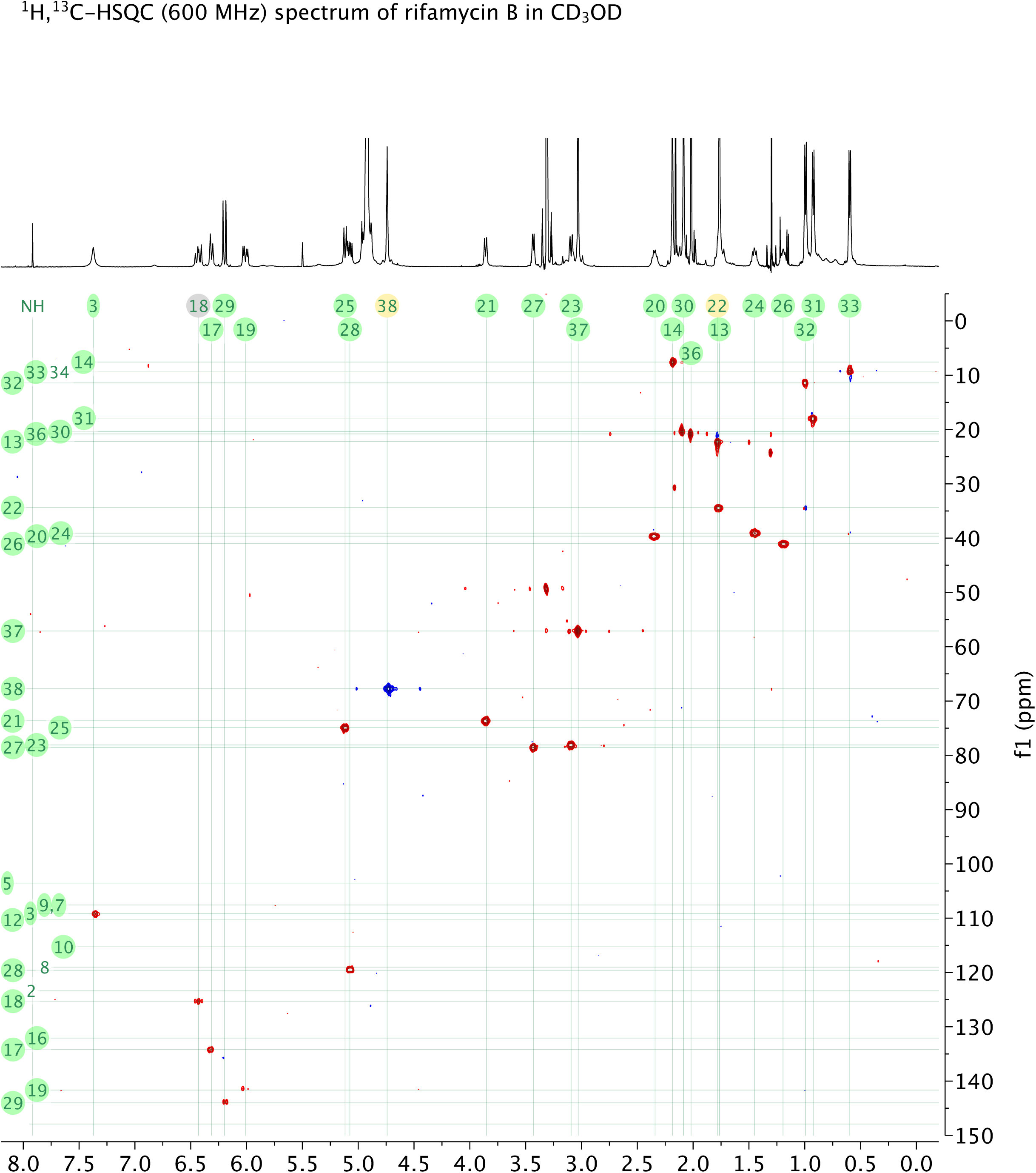
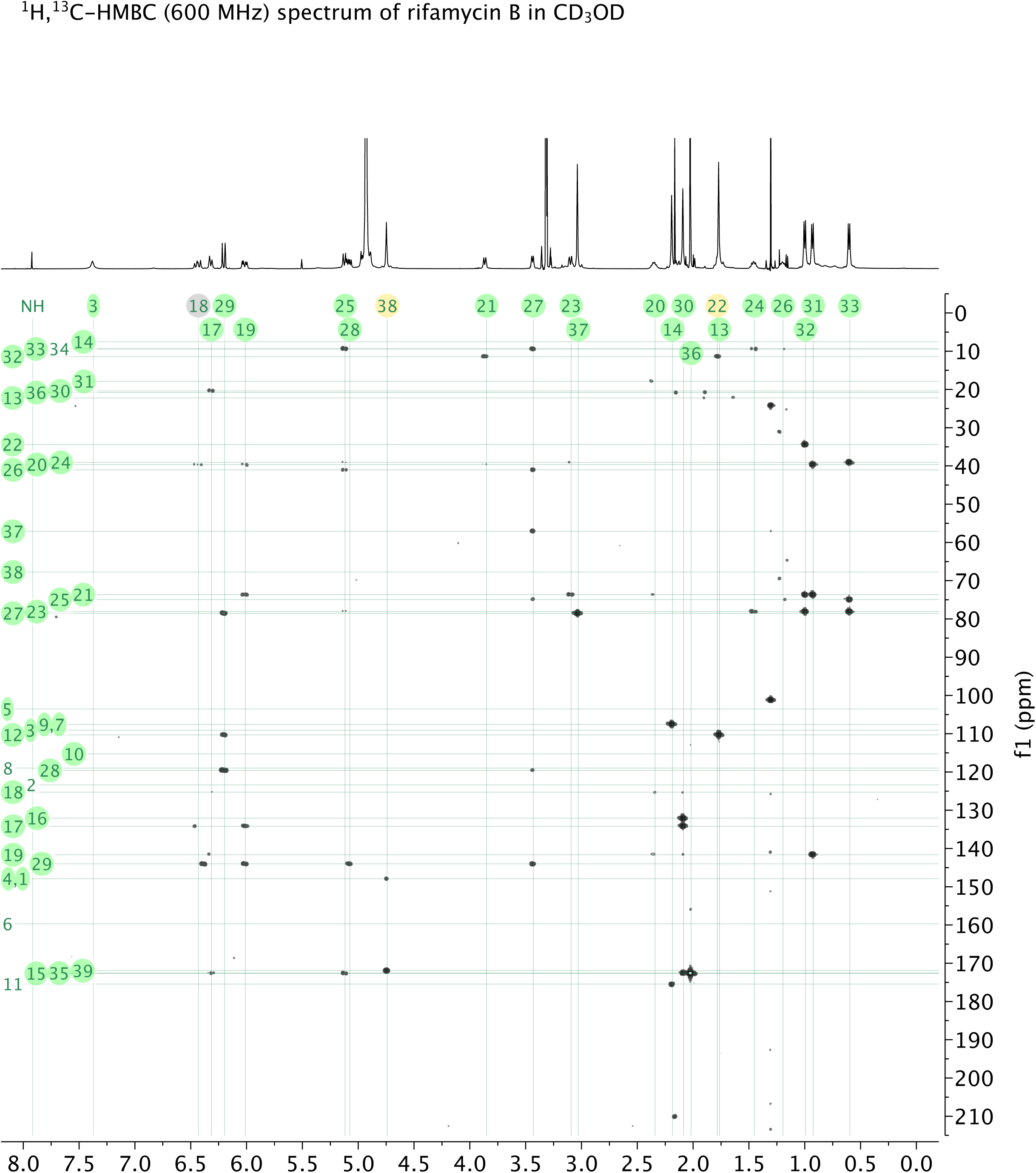
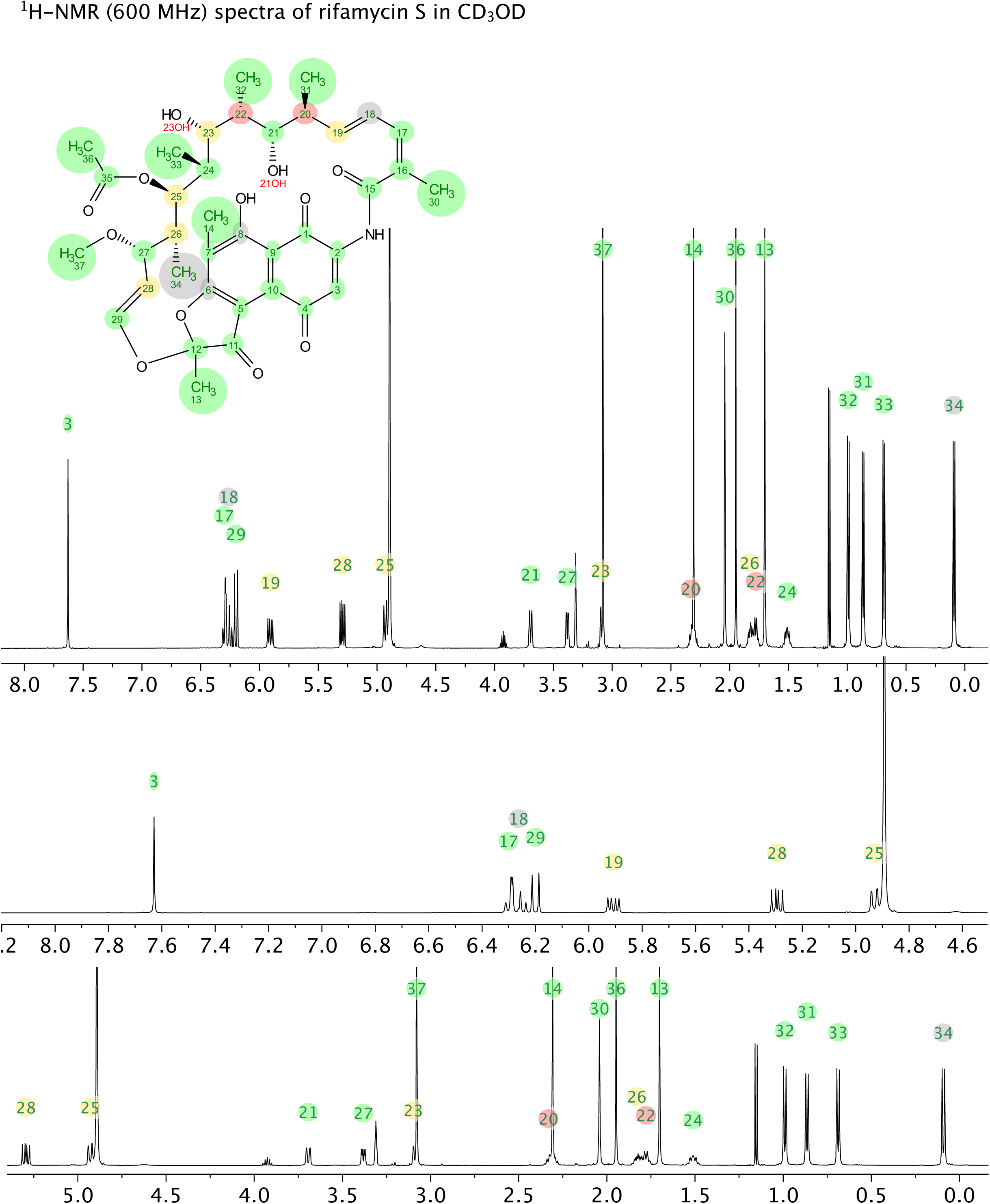
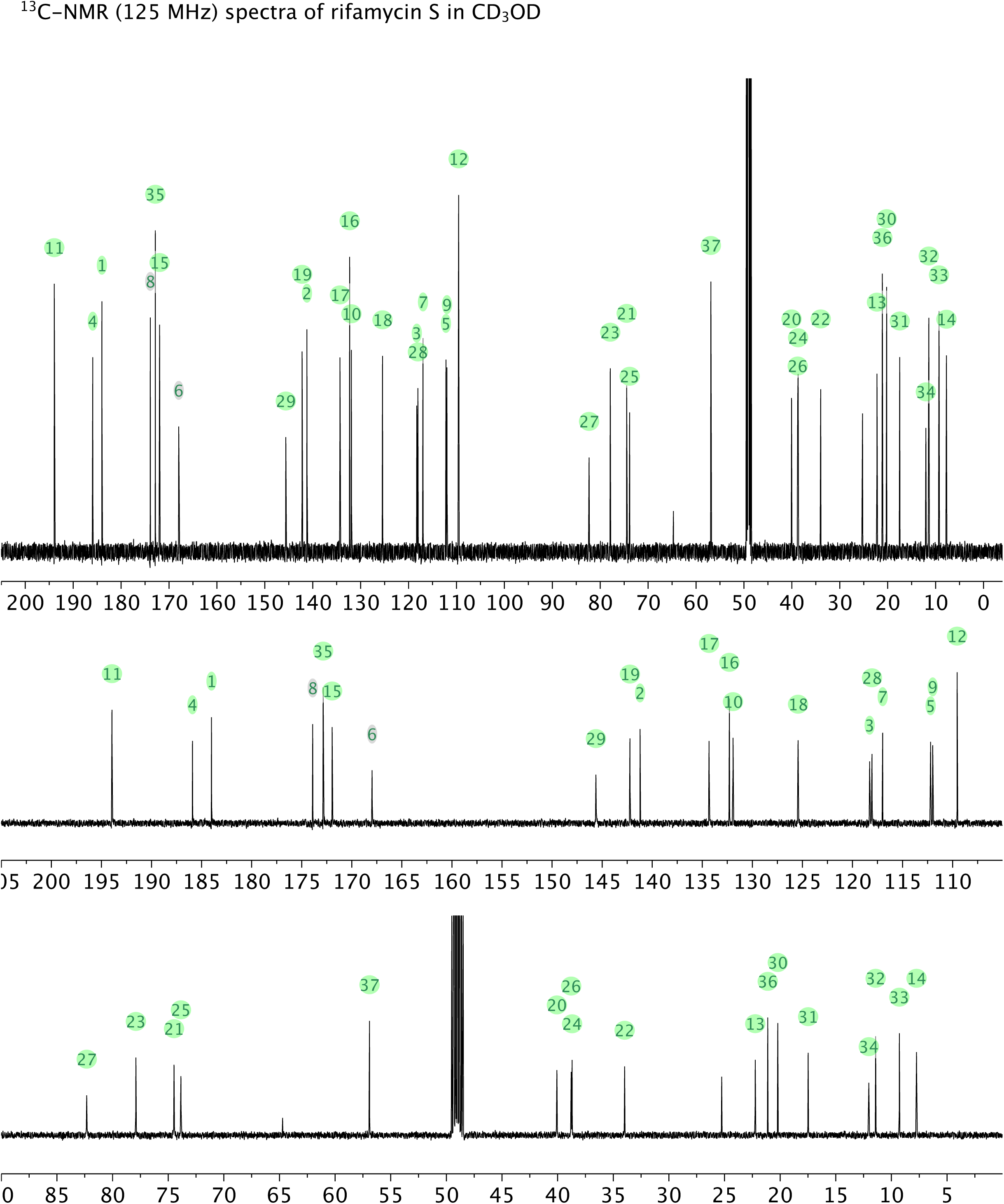
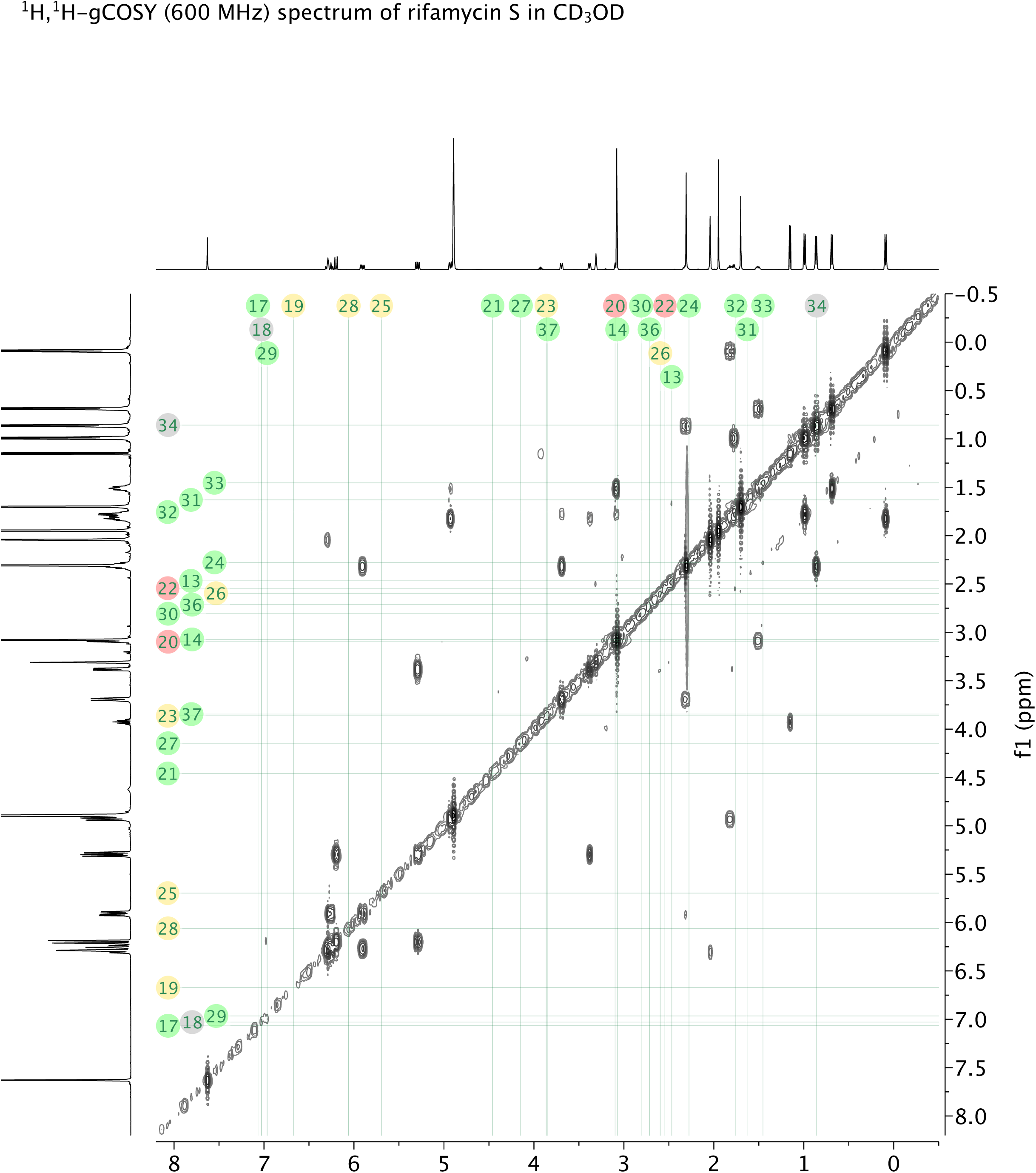
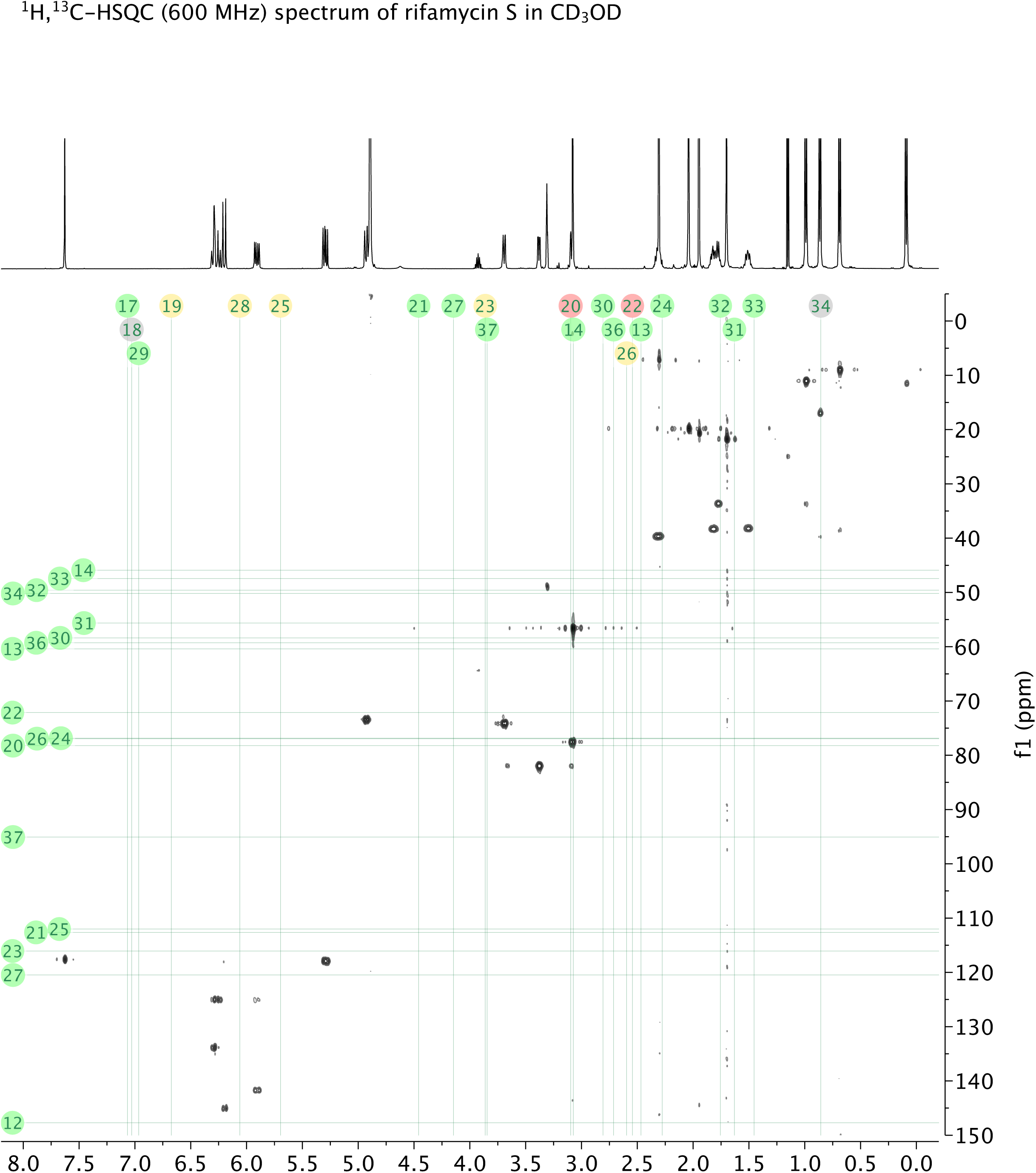
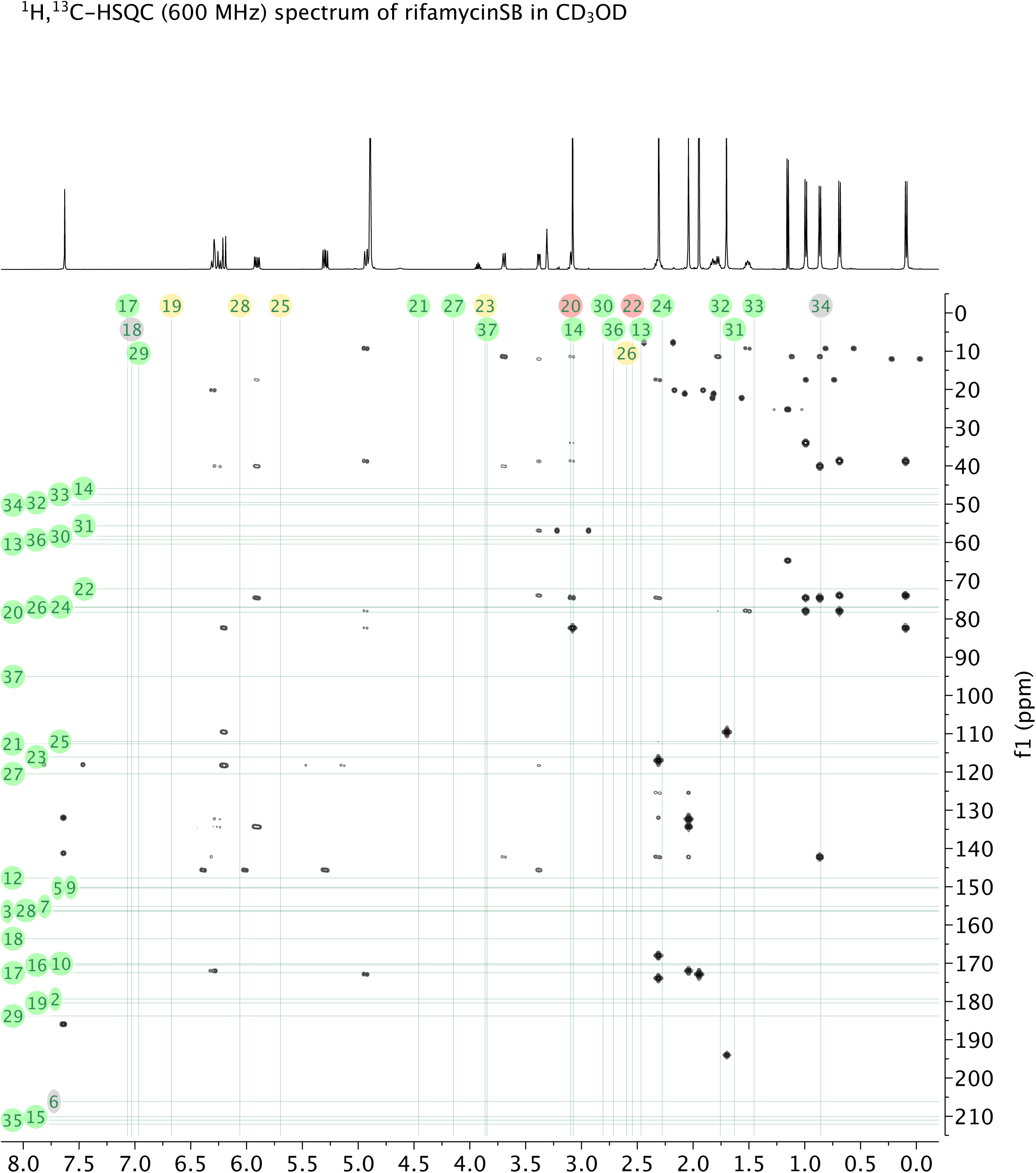

